# Haemodynamic impact of implant materials and anastomotic angle in peripheral vascular grafts

**DOI:** 10.1101/2024.12.18.629298

**Authors:** Sabrina Schoenborn, Thomas Lloyd, Yogeesan Sivakumaran, Maria A. Woodruff, David F. Fletcher, Selene Pirola, Mark C. Allenby

## Abstract

End-to-side anastomoses are commonly utilised in peripheral arterial bypass surgery and are plagued by high rates of re-stenosis as a result of non-physiological blood flow impacting arterial and graft structures. Computational simulations can examine how patient-specific surgical decisions in bypass graft placement and material selection affect blood flow and future risk of graft restenosis. Despite graft geometry and compliance being key predictors of restenosis, current simulations do not consider the interaction of flowing blood with compliant vessel, graft, and suture structures. Utilising fluid-structure interaction simulations, this study examines the impact of surgical technique, such as anastomosis angle, graft material, and suture material, on blood flow and fluid-structure forces in patient-specific asymptomatic arterial tree versus side-to-end peripheral grafts for symptomatic atherosclerotic disease. To render these complex simulations numerically feasible, our pipeline uses regional suture mechanics and a pre-stress pipeline previously validated in small-scale idealised models. Our simulations found that higher anastomosis angles generate larger regions of slow and recirculating blood, characterised by non-physiologically low shear stress and high oscillatory shear index. The use of compliant graft materials reduces regions of non-physiologically high shear stress only when used in combination with compliant suture materials. Altogether, our fluid-structure interaction simulation provides patient-specific platforms for vascular surgery decisions concerning graft geometry and material.

**Highlights:**

- Simulating bypass graft haemodynamics with realistic fluid-structure interactions.
- Bypass grafts generate large regions of slow blood flow and blood recirculation.
- Greater graft anastomosis angles correlate with larger blood recirculation regions.
- Nonphysiologically stiff graft and suture materials increase vessel shear stress.

## 1. Introduction

Cardiovascular disease (CVD) is the most common cause of death worldwide. The annual incidence of global deaths related to CVD is expected to rise from 16.7 million in 2002 to 23.3 million in 2030 [1]. A common manifestation of CVD is peripheral arterial disease (PAD), whose symptoms develop as a consequence of the critical atherosclerotic narrowing of the peripheral arteries. International research studies show a prevalence of approximately 10–20% for PAD depending on study design, country, and participant’s age and gender [2]. PAD affects more than 200 million people worldwide, with atherosclerosis accounting for more than 90% of cases of PAD. This study focuses on the femoro-popliteal segment, since it is the arterial segment with the highest likelihood to develop PAD [3,4] with the femoral and popliteal arteries being affected in 80–90% of symptomatic PAD patients [5].

In the case of PAD progression to critical limb threatening ischaemia (CLTI), restoration of tissue perfusion is essential for limb salvage. Current revascularisation techniques consist of endovascular interventions, including angioplasty, stenting and other endovascular adjuncts (atherectomy and catheter directed thrombolysis), as well as open surgical repair with bypass grafting using autologous or prosthetic conduit. The choice of operative treatment for CLTI has been evaluated recently in the BASIL-2 and Best-CLI trials which demonstrated disparate results when comparing open versus endovascular revascularisation techniques for CLTI [6,7]. From the perspective of lower limb bypass surgery, the gold standard conduit is autologous, with the great saphenous vein demonstrating the best patency rates at all intervals and the fewest re-interventions [8]. Great saphenous vein as a bypass graft conduit has an average primary patency rates of 74% after 5 years [9]. However, the use of autologous grafts require harvesting and may be unavailable for use as a result of prior bypass grafting procedures or pre-existing conditions (varicose degeneration) [10]. One-third of patients requiring primary revascularisation lack a suitable autologous conduit. This number increases to 50% in patients presenting for a re-operation or secondary bypass procedure [11].

In cases where no autologous graft can be used, blood flow can be restored by implanting synthetic grafts. The two most common materials for synthetic vascular grafts are expanded poly(tetrafluoroethylene) (ePTFE) and poly(ethylene terephthalate) (PET), better known by its brand name Dacron^®^ [12]. While synthetic grafts are readily accessible in all required lengths and diameters, their low average primary patency rates of 39% after 5 years are their most significant limitation [9]. The economic burden associated with PAD is considerable because of the high rates of leg revascularisation procedures and hospitalisations [13] and the poorer results of secondary bypass surgeries that may be required after primary bypass graft failure [14].

While early graft failure within 30 days of bypass grafting is usually due to complications at the time of surgery, such as technical errors, graft thrombogenicity, low flow, or poor run-off, late graft failure is mainly caused by the graft abnormality and metachronous disease. In particular, graft stenoses due to intimal hyperplasia [15–18] at points of anastomosis have been reported as a major cause of failure in autogenous and prosthetic bypass [10,18–20].

Intimal hyperplasia (IH) is the chronic and excessive thickening of the tunica intima, caused by abnormal proliferation and migration of vascular smooth muscle cells (VSMCs) in response to endothelial injury or dysfunction with associated deposition of connective extracellular matrix (ECM), which results in either partial or complete occlusion of the blood vessel [21–23]. IH leads to graft failure due to restenosis in approximately 20-30% of femoro-distal bypasses utilising autologous conduit and is recognised as the main cause of thrombotic complications occurring between 2 and 24 months after a vascular intervention [24]. Current surgical anastomotic interventions for vascular anastomoses utilise a monofilament suture, with a continuous polypropylene (Prolene^®^) suture being considered the gold standard. Regardless of the choice of suture material and technique, suturing results in injury of the vessel wall and the presence of foreign material at the blood-graft interface [22].

A range of biological, chemical, and mechanical factors have been studied with regards to their potential to aggravate the severity of IH [25], and compliance mismatch of graft and anastomosis with the host artery is one of the central biomechanical issues of vascular graft design, as greater compliance mismatch is associated with greater incidence and severity of IH and consequently lower mid-term patency rates [24,26].

Combined in-vivo and in-silico studies on small-diameter arterial models have demonstrated that the severity of IH is related to local biomechanical factors. In particular, regions most significantly affected by IH are the toe, heel, and floor of the anastomosis [27,28]. The sudden change in flow direction at the anastomosis site induces regions of both high and low wall shear stress (WSS) and disturbed flow profiles, including secondary, recirculating, and oscillatory flows [18]. Both the magnitude and the level of flow disturbance as marked by the oscillatory shear index have been indicated to affect IH severity [29]. Additionally, high intramural stresses at the suture line due to the presence of stiff graft and suture materials have been indicated to have a proliferative effect on IH around the suture [30], with the degree of IH linked to the mechanical compliance mismatch between the connected materials [26].

Fluid-structure interaction (FSI) simulations are particularly useful where changes in vascular compliance, which may occur during disease or after surgical intervention, are of interest. Examples of such applications are studies on the increase in stroke work after aortic aneurysm repair with a stiff graft [31], the effect of graft versus stent repair on haemodynamics [32], vascular wall stress profiles after aortic grafting [33], the identification of adverse oscillatory flow after implantation of stiff Dacron^®^ grafts [34], the investigation of suitable graft materials for aortic repair [35], the effect of different surgical orientations and graft length ratios on haemodynamics [36].

Due to the complexity of the biomechanical causes of IH encompassing both the blood flow and the arterial wall, this study aims to analyse both the geometric factors, such as anastomosis angle, and the structural factor, such as choice of suture and graft material, independently utilising a multi-patient FSI study.

## 2. Computational Methods

A FSI model coupling blood flow with artery, graft, and suture wall motion was developed using the 2023R1 version of the commercial computational software suite Ansys (Ansys Inc., Pennsylvania). Ansys Mechanical was used for the structural mechanics analysis and Ansys CFX was used for the fluid mechanics computations.

### 2.1 Image Segmentation and Meshing

Patient-specific asymptomatic arterial disease (***n*** = ***5***) and bypass graft (***n*** = ***5***) vascular geometries were obtained from computed tomography angiography (CTA) image datasets [37]. The 3D imaging software Materialise Mimics was used to manually segment the contrast-enhanced blood model using thresholding, inter-slice interpolation, region growing tools, and smoothing tools [38]. Volume and shape conserving algorithms were used. Cross-sectional areas of graft, anastomosis and artery were checked pre- and post-smoothing, and less than 2% deviation was achieved. The model was exported in STL format and further processed in Autodesk Meshmixer. Since the imaging data included all vascular structures distal to the diaphragm, the vascular model was cut to the region of interest around the distal anastomosis while maintaining minimum inflow and outflow lengths [39]. The structural models were generated from the segmented fluid geometry via an offset function of 1.2 mm [40] and 0.64 mm [41,42] for the arterial and graft segments, respectively. The latter was assumed to be representative of the thickness of both vein and ePTFE grafts. Both structural and fluid models were meshed in Ansys ICEM [43,44] as unstructured tetrahedral meshes with an average size of 0.3 mm [45]. Ten boundary fitted prism layers decreasing in size towards the wall were generated to ensure more accurate WSS data [46], as shown in **Figure 1**. A transient mesh sensitivity analysis was performed for each patient case with regards to velocity, pressure, and wall shear stress for the fluid mesh, and deformation and von-Mises stress for the structural mesh. The mesh study was conducted on vein grafts with polypropylene sutures, since this material combination led to the highest von-Mises stresses. Results were considered mesh-independent when the difference of these parameters was less than 2% compared with a mesh with an increase of 150% in the number of nodes. Full details of the conducted sensitivity tests can be found in the supplementary materials.

**Figure 1:**
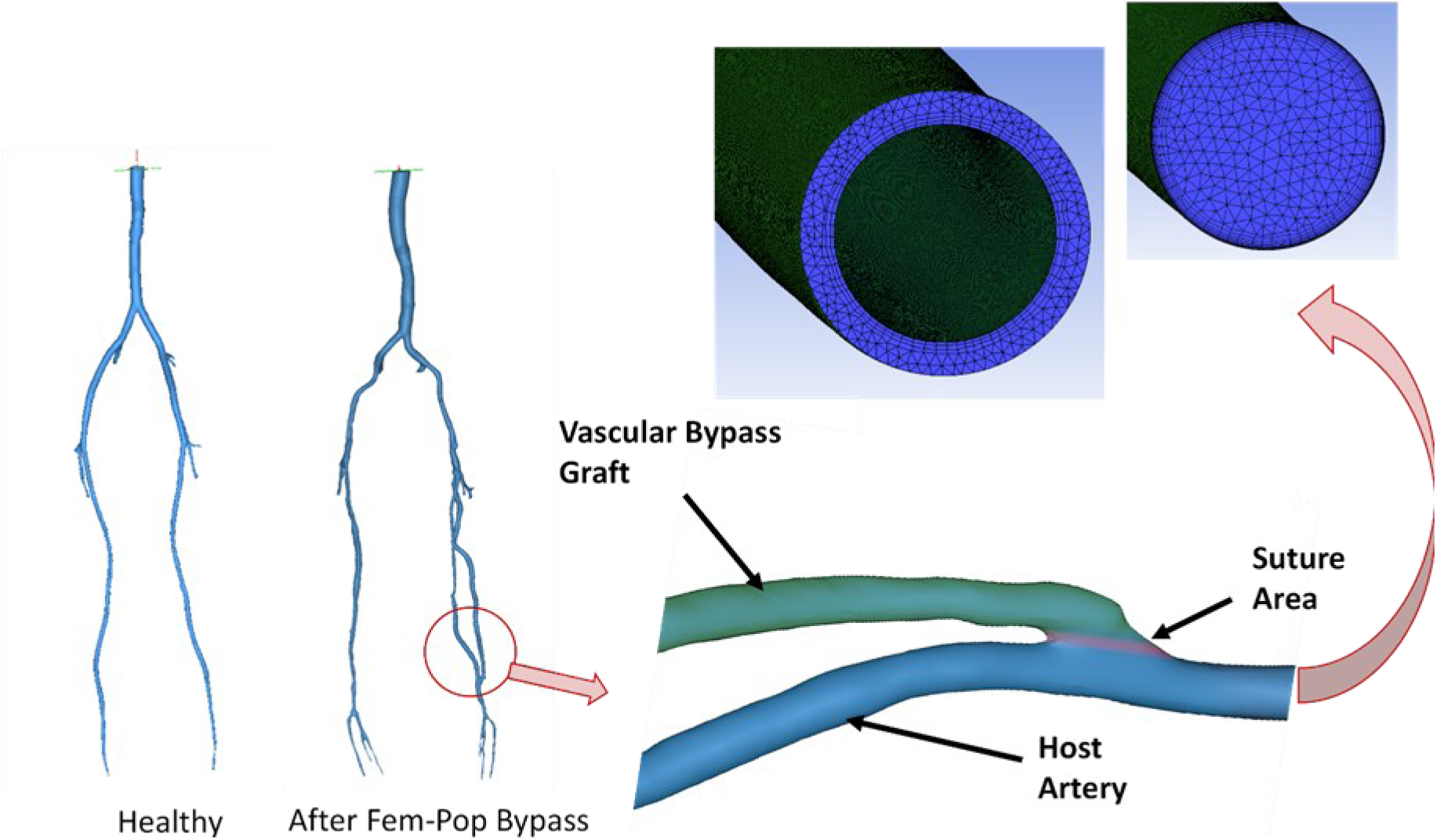
Structural model generation from STL geometry to material assignment to meshing.

### 2.2 Blood Fluid Model

In this study, the Carreau-Yasuda model was used to model the blood viscosity [46,47]. The apparent viscosity *η* is given by the shear rate 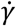, empirically determined constant parameters *a*, *n*, and *λ*, and viscosities 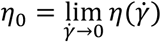 and 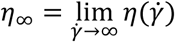

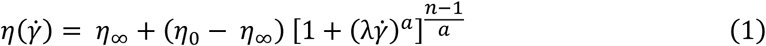

with *η*_∞_ = 0.0035 Pa·s, *η*_0_ = 0.1600 Pa·s, *λ* = 8.2 s, *a* = 0.64, and *n* = 0.2128 [47].

A time-dependent velocity profile was prescribed at the inlet of the fluid model as the inflow boundary condition and a time-dependent pressure profile was prescribed at the outlet. Boundary conditions were assumed to be uniform over the cross-section of the inlet and outlet. Both the inflow and outflow velocity or pressure boundary conditions were described as eighth-order Fourier series fit:

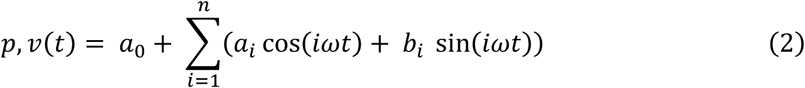

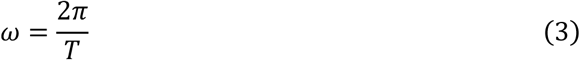

where *a*_0_ models a constant intercept term in the data and is associated with the *i* = 0 cosine term, *ω* is the fundamental frequency of the signal, *T* is the signal’s period, i.e., the cardiac cycle length, and *n* (= 8) is the number of terms in the series. The Fourier series were fitted to pressure and velocity data taken from the Pulse Wave Database (PWD) for a 65-year-old patient [48]. The model was fitted for two heartbeats to ensure periodicity, using the MATLAB (MathWorks, Massachusetts) Curve Fitting tool (**Table S1**).

**Table 1:** First-order Ogden model parameters of structural arterial wall model. Incompressibility parameter d_1_, material constant α_1_, material constant μ_1_, density ρ.

| Parameter | Value | Source [Reference] |
| --- | --- | --- |
| $\mu_1$ | 26.414 kPa | Ansys curve fitting tool, source data from literature [56] |
| $\alpha_1$ | 14.405 | Ansys curve fitting tool, source data from literature [56] |
| $d_1$ | $3.63 \times 10^{-6} \text{ Pa}^{-1}$ | Literature [55] |
| $\rho$ | $1080 \text{ kg m}^{-3}$ | Literature [60] |

A transient analysis was performed with a Second Order Backward Euler transient scheme and high resolution (2nd order iteratively bounded) advection scheme. In asymptomatic patient models, a laminar fluid model was applied with a time step size of 10^−3^ s. This was selected via a timestep sensitivity analysis. Timestep convergence was considered achieved when the change in wall shear stress, velocity, and pressure was less than 1% for a given timepoint after halving the time step size (please refer to supplementary materials). In patient models with bypass grafts, laminar simulations were compared with those using scale-resolving turbulence models such as the Stress Blended Eddy Simulation (SBES) model, which applies the *k*-*ω* SST model in the near-wall region and a Large Eddy Simulation model away from the wall with the transition between the two accomplished using a blending function. No turbulent flow features were found, which is in agreement with typical Reynolds numbers in peripheral arteries and the current literature [49]. A timestep was considered as converged when the Root Mean Square (RMS) residuals of the mass conservation equation and the momentum conservation equations were all below a residual target of 10^−4^ and the mesh deformation equation was below a residual target of 10^−5^. The model was run for two cardiac cycles after confirmation of cyclic convergence, where differences in wall shear stress, velocity, and pressure between cycle 2 and 3 were less than 0.5% for each corresponding timestep, and only the last cycle was analysed to eliminate transient start-up effects from the results.

Since flow properties change throughout the cardiac cycle, time-averaged parameters such as the time-averaged WSS magnitude (TAWSS) and time-averaged WSS vector (TAWSSV) are commonly analysed in biological flows [50]:

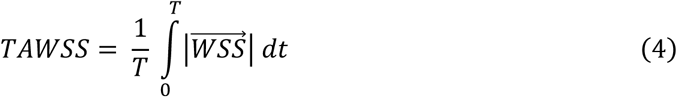

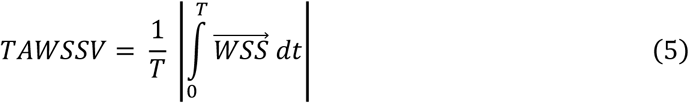

Both parameters can be combined into the oscillatory shear index (OSI), with OSI values larger than 0.15 being considered a risk factor in the development of arteriosclerosis and IH [51,52] since they indicate large temporal variations of low and high WSS being present in recirculation zones:

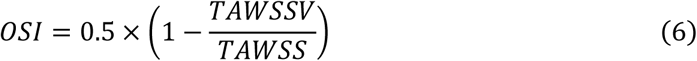

### 2.3 Artery Structural Model

A hyperelastic first-order Ogden model was utilised to reproduce the arterial wall’s stress-strain behaviour [53–55]. The Ogden model parameters were determined using the Ansys hyperelastic curve fitting function on circumferential uniaxial tensile testing data of healthy femoral arteries reported in the literature [56]. The model parameters utilised for a 65-year-old patient are calculated via the Ansys Curve Fitting Tool for hyperelastic materials using published mechanical testing data on femoral arteries and are summarised in **Table 1**. The tissue surrounding the femoral artery was expected to have a damping effect on the arterial wall movement, which was taken into account by applying a linear elastic support boundary condition [57]. The structural model was run as a transient analysis using the sparse direct solver with a time-step size matching the fluid solver. A timestep was considered as converged when the force convergence value and the displacement convergence value were both below a residual target of 10^−4^. The structural model was pre-stressed using a ten-step iterative pre-stress pipeline following Caimi *et al.* [58], which we previously adapted for Ansys and experimentally validated in idealised small-diameter cylindrical models [59].

### 2.4 Graft and Suture Structural Models

Idealised models of artery-graft anastomoses explicitly modelling the sutures are employed to accurately model the stresses and deformations at the anastomosis region for different graft-suture material combinations. The results from these idealised models were utilised to inform isotropic linearly elastic material models to model the anastomotic region in the patient-specific simulations (**Figure 1**). Specifically, the isotropic linear elastic material parameters in the simplified models were calibrated to match the deformation and von-Mises stress responses of the explicitly modelled sutures in the idealised models within a 2% tolerance averaged across the stitch length. The rationale behind this approach is primarily driven by computational efficiency as the inclusion of suture contacts in full-scale 3D patient-specific FSI simulations would increase the computational cost for each patient case to impractical magnitudes. This method effectively strikes a balance between the analysis of suture line stresses without significant loss of accuracy with the possibility to study a larger number of patient models and follows similar strategies employed in other numerical patient-specific vascular models in literature, particularly in the study of stents and grafts [61].

A continuous suture with a circular cross-section and diameter *D*_O,Suture_ = 0.07 mm [62] was modelled as depicted in **Figure 2**. The contact between suture and artery or graft is assumed to be a frictional contact with a coefficient of friction *μ_F_* = 0.43.

**Figure 2:**
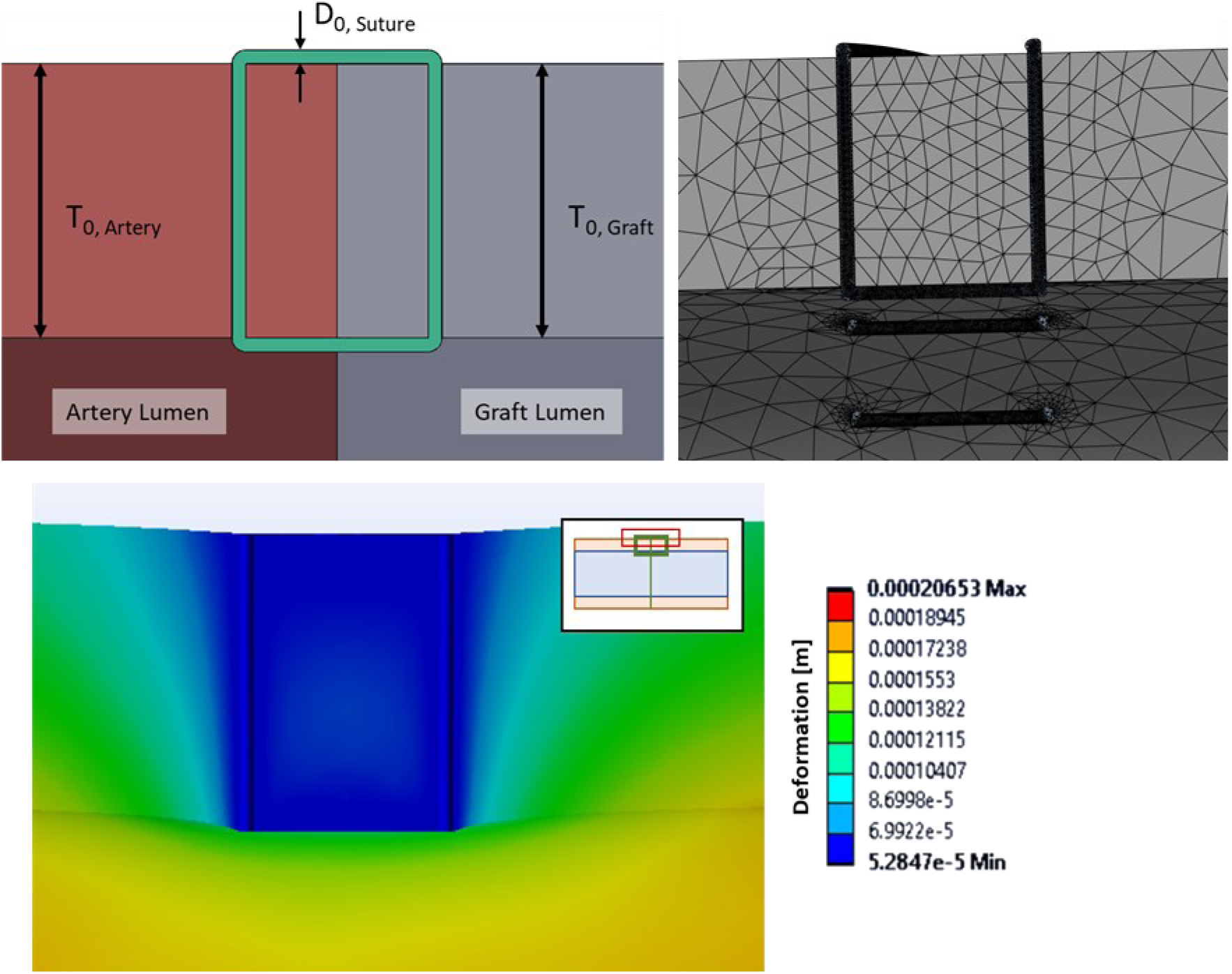
Simplified geometry of a continuous polypropylene suture with a diameter D_O,Suture_ in an anastomosis between an artery with thickness T_O,Artery_ and graft with thickness T_O,Graft_.

The elastic modulus of the PP suture is determined by linear regression using tensile testing data published by Dobrin *et al.* [63] and is in agreement with data published on the mechanical properties of PP elsewhere [64]. The suture materials are modelled as linearly elastic as outlined in **Table 2**. The PP suture is compared with an alternative more compliant suture made of polybutester (PB).

**Table 2:** Properties of the polypropylene and polybutester sutures.

| Material | Density [kg/m<sup>3</sup>] | Elastic Modulus [MPa] | Poisson's Ratio |
| --- | --- | --- | --- |
| Polypropylene | 900 [64] | 1483 [63,64] | 0.42 [64] |
| Polybutester | 1220 [65] | 196.2 [66] | 0.35 [67] |

Similar to the femoral arterial wall, the saphenous veinous wall shows a J-shaped nonlinear mechanical behaviour. While the saphenous vein is more compliant than the femoral artery at the low luminal pressures typically present in the venous system, it is markedly less circumferentially compliant at the high luminal pressures it is subjected to after implantation into the arterial system [68,69]. This nonlinear stiffening of the veinous wall leads to a compliance mismatch with the more compliant arterial wall under arterial pressure, which is however less pronounced than the compliance mismatch between artery and synthetic graft materials, such as ePTFE [69]. A first-order Ogden model was chosen to describe the behaviour of the saphenous veinous wall graft material. The model parameters utilised for saphenous vein graft are summarised in **Table 3**. A linear elastic material model was chosen to describe the behaviour of the ePTFE graft material (**Table 4**).

**Table 3:** First-order Ogden model parameters of structural saphenous vein graft model. Incompressibility parameter d_1_, material constant α_1_, material constant μ_1_, density ρ.

| Parameter | Value | Source [Reference] |
| --- | --- | --- |
| $\mu_1$ | 8.577 kPa | Ansys curve fitting tool, source data from literature [70] |
| $\alpha_1$ | 65.221 | Ansys curve fitting tool, source data from literature [70] |
| $d_1$ | $3.63 \times 10^{-6} \text{ Pa}^{-1}$ | Literature [55] |
| $\rho$ | $1080 \text{ kg m}^{-3}$ | Literature [60] |

**Table 4:**
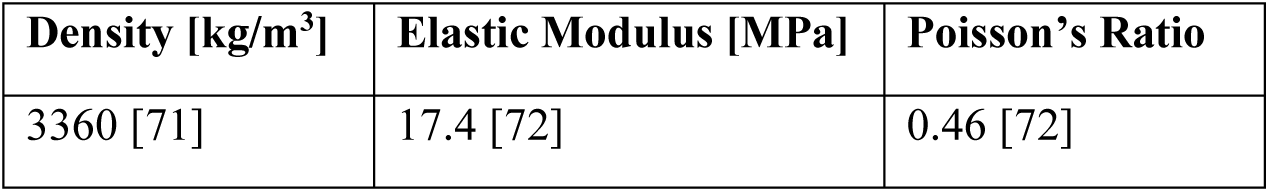
Properties of the ePTFE graft model.

### 2.5 Fluid-Structure Interaction System Coupling

Ansys System Coupling was used to facilitate two-way data transfers between the outer surface of the fluid volume and on the luminal surface of the structural model [73]. A time step was considered converged when the individual solvers had converged and the force and displacement coupling data transfers had met an RMS convergence target of 10^−2^ [74,75]. Fluid mesh deformation was calculated using a displacement diffusion model without remeshing. The flow solution was stabilised to prevent divergence by setting a velocity-dependent source coefficient in the pressure equation at the arterial wall and monitoring monotonical convergence for a given time step by plotting the forces and displacements at the arterial wall boundary [76].

### 2.6 Statistical Analysis

Statistically-significant differences between simulated asymptomatic and bypass graft patient anastomosis angles, percent area of high oscillatory shear index (OSI > 0.15), and percent area of high or low time-averaged wall shear stress (TAWSS > 3 Pa or TAWSS < 0.5 Pa) were calculated using *p*-values from Student’s bidirectional *t*-test, assuming unpaired asymptomatic and grafted patients of equal variance. Linear correlations between anastomosis angle with percent area of high oscillatory shear index, or high, or low time-averaged wall shear stress were calculated from Spearman’s R^2^ coefficient, as shown in **Figure 6**.

## 3. Results

### 3.1 Greater Anastomotic Angles Enlarge ‘Toe’ Regions of Low Blood Shear Stress and High Oscillatory Blood Flow

Proximal to the anastomosis, a fully developed laminar Womersley flow profile through the graft is observed with no flow in the occluded artery. At the anastomosis, a transition to disturbed flow is observed with high velocity and WSS at the toe of the anastomosis and eddies and dead zones at the heel of the anastomosis (**Figure 3**). At and distal to the anastomosis, zones of recirculation and oscillatory flow behaviour were observed in all five patient cases, resulting in 25.29% to 72.59% of regions around the anastomosis (± 5 cm).

**Figure 3:**
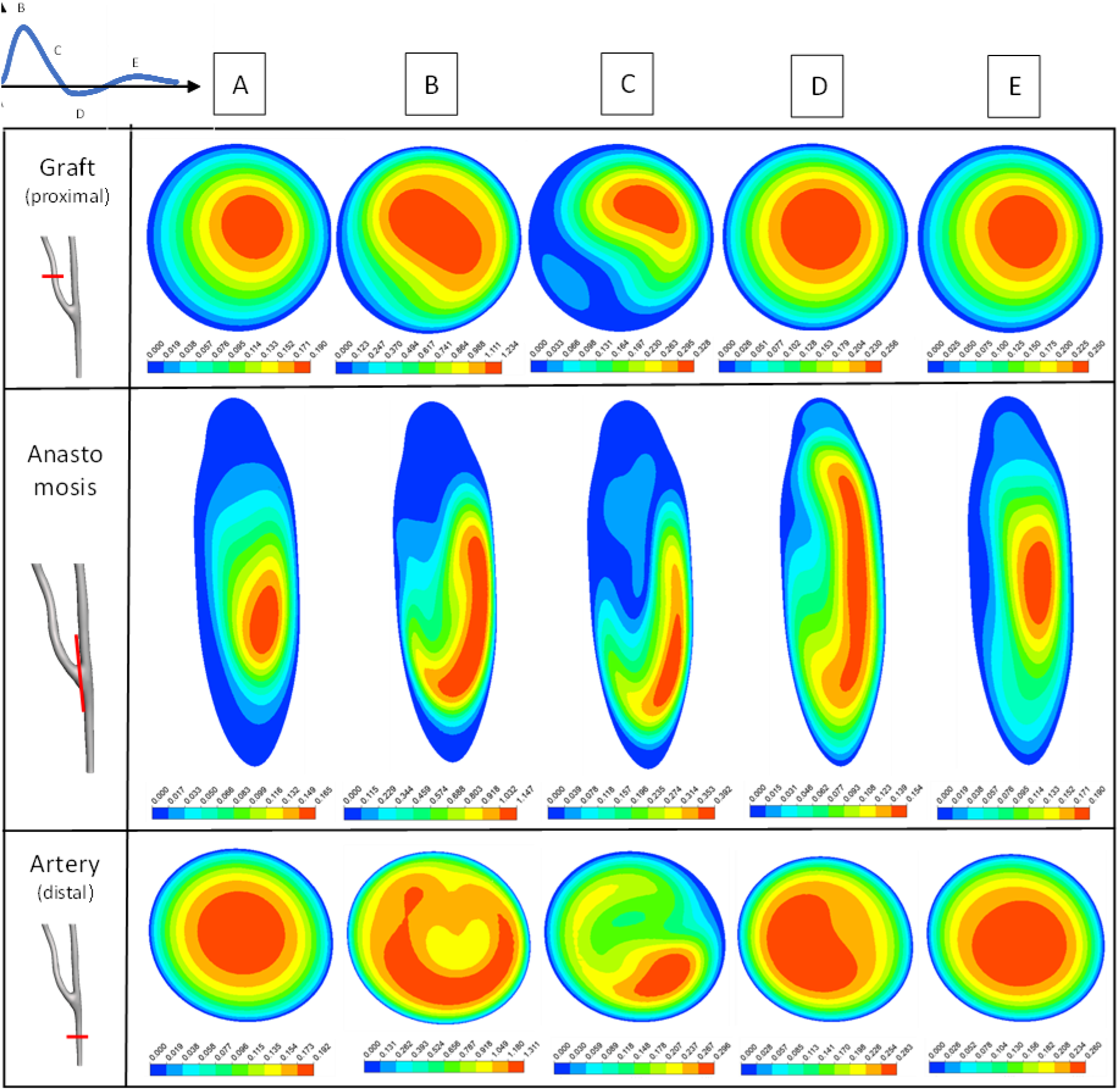
Local velocity magnitude (in m·s^−1^) profiles in femoral-popliteal bypass graft (top) proximal to distal anastomosis, (middle) at distal anastomosis, and (bottom) in host artery distal to distal anastomosis along cardiac cycle. Each of these three locations contains five cross-sectional velocity magnitude profiles, corresponding to five time points along a cardiac cycle (as illustrated top-left).

In all bypass graft patients, a recirculation zone is observed at the toe of the anastomosis, with the size of the recirculation zone being positively correlated with larger anastomosis angles (R^2^ = 0.96, **Figure 6B**) and appearing to have larger-magnitude downstream vorticity, consistent with prior experimental flow models [77]. A larger, i.e., more obtuse anastomosis angle as measured between the graft and host artery towards the heel of the anastomosis, is associated with a larger region of the arterial wall subjected to high OSI and low TAWSS values (TAWSS < 0.5 Pa; **Figure 6** and **Table S2**). Conversely, the lowest anastomosis angles were slightly correlated to have larger regions experiencing high TAWSS values (TAWSS > 3 Pa; R^2^=0.56; **Figure 6C**).

In the asymptomatic femoral-popliteal segments, regions of disturbed flow (OSI > 0.15) were significantly reduced compared with the grafted patient cases with 11.92% ± 7.81% vs 50.16% ± 15.93% (*p* = 0.005) (Figure 6). There is no statistically significant difference in the areas of high WSS (TAWSS > 3 Pa) with 2.55% ± 2.00% for asymptomatic vs 3.41% ± 5.26% for grafted patient cases (*p* = 0.8) (Figure 6). Conversely, there were no areas of low WSS (TAWSS < 0.5 Pa) in the asymptomatic geometries, vs 10.56 ± 2.90 % in the grafted geometries (*p* = 0.002) as shown in **Figure 5** and **Table S3**.

**Figure 4:**
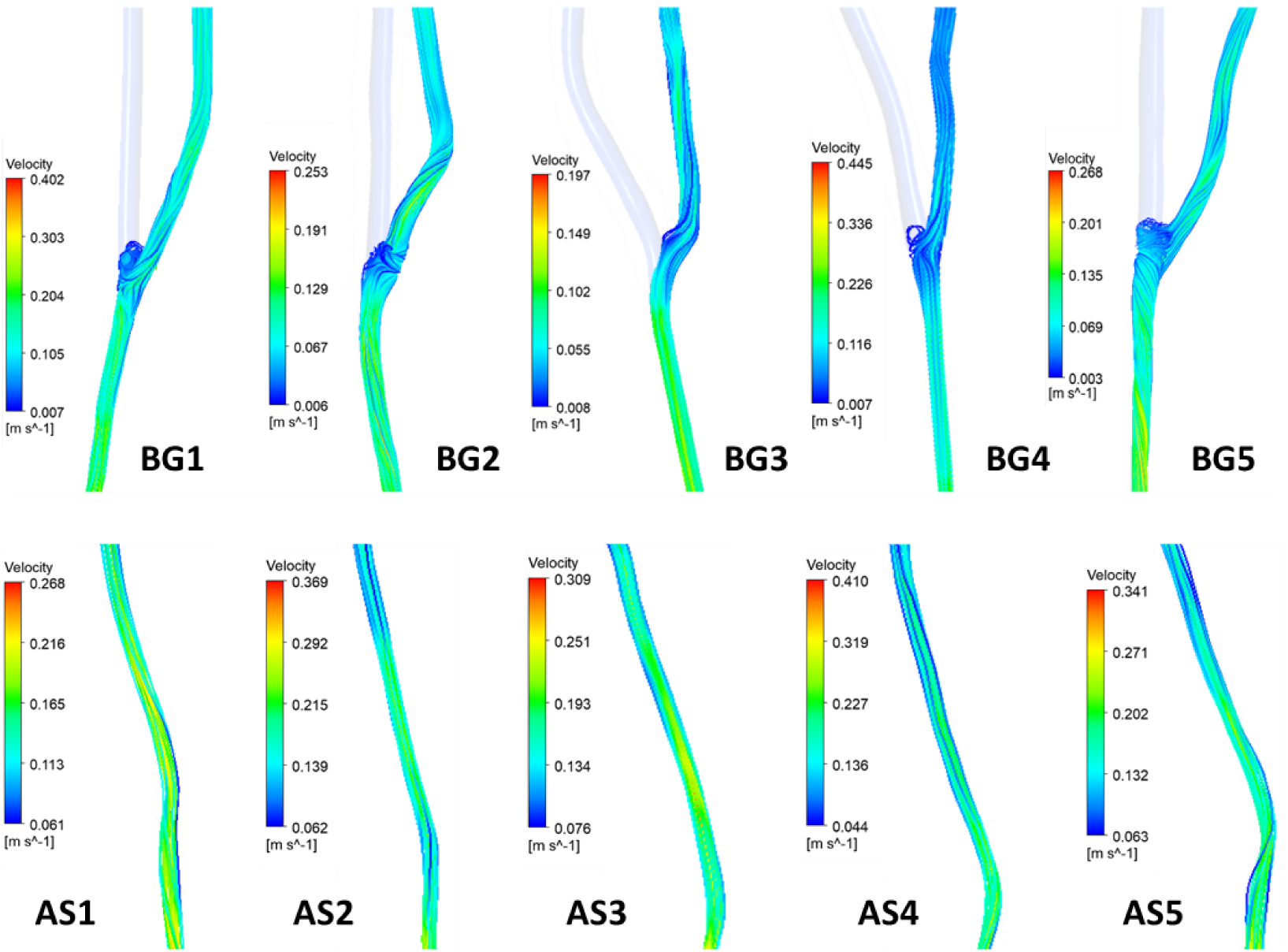
Velocity streaklines at mid-diastole for five patient-specific ePTFE-PP distal femoral-popliteal bypass graft models (BG; top row) and for five patient-specific asymptomatic femoral-popliteal segment models (AS; bottom row).

**Figure 5:**
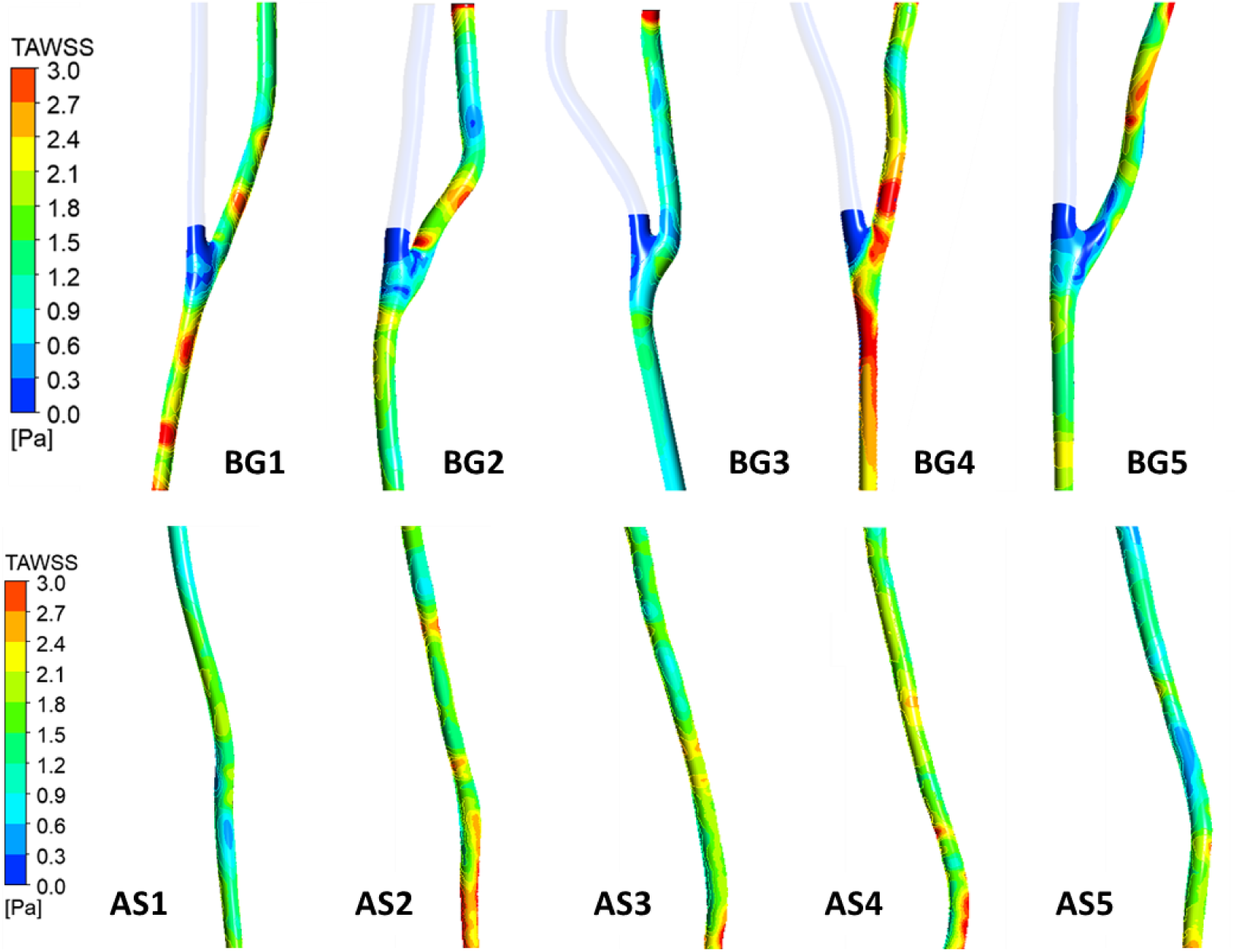
Time-averaged wall shear stresses for patient-specific ePTFE-PP distal femoral-popliteal bypass graft models (BG; top row) and of patient-specific asymptomatic femoral-popliteal segment models (AS; bottom row).

**Figure 6:**
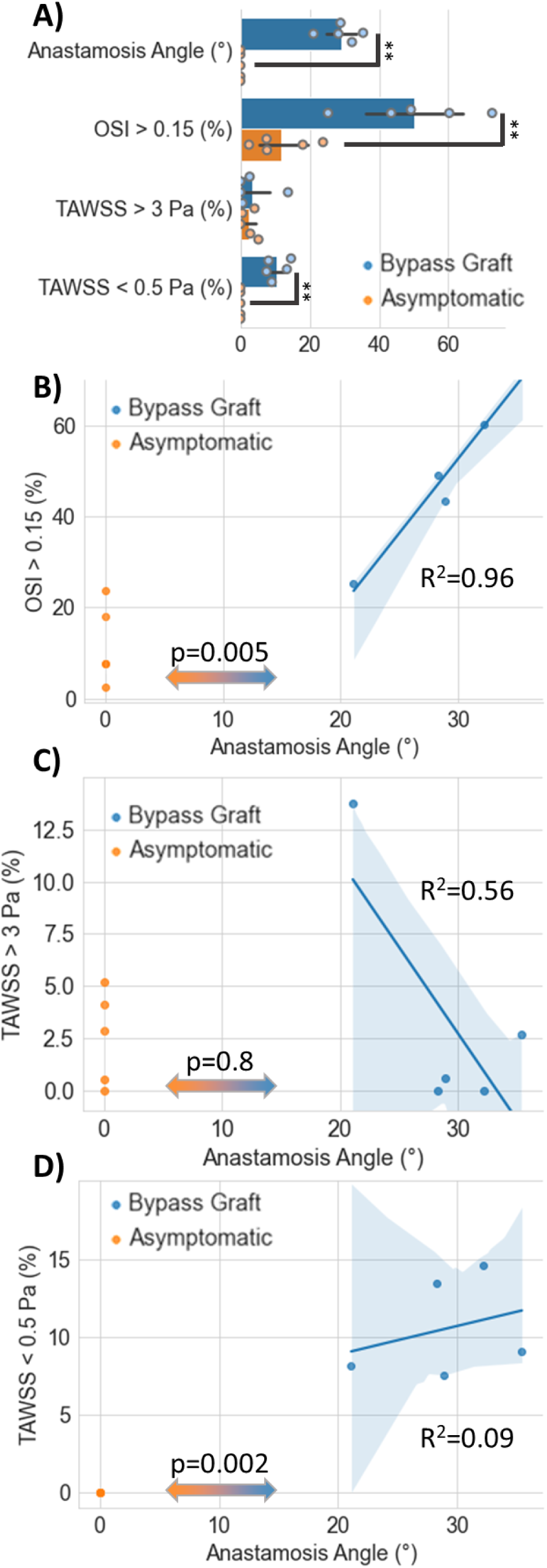
Summary plots comparing simulations for ePTFE-PP grafted patients (in blue) and asymptomatic controls (in orange). Pairwise student’s t-tests were preformed between asymptomatic and grafted patient scans, with p-values annotated in (A) and then repeated in (B, C, D). Pearson’s R^2^-coefficient was applied to measure linear correlations between graft anastomosis angle and OSI or TAWSS percent. Exact values for anastamosis angle, OSI shear index, or TAWSS percentage can be found for each patient in **Tables S2, S3.**

### 3.2 Stiffer Graft and Suture Materials lead to Larger Regions of High Stress and Strain

The compliance of graft and suture showed small effects on tested fluid parameters for bypass graft patient BG1. For this single patient replicate, regions of elevated OSI made up 72.59%, 72.46%, 72.39%, and 72.69% of ePTFE-PP, ePTFE-PB, Venous-PP, and Venous-PB anastomotic regions, respectively. For patient BG1, TAWSS were reduced for the more compliant saphenous vein graft material with anastomotic regions being subjected to areas of 2.70%, 2.73%, 0.49%, and 0.31% of high TAWSS and 9.06%, 9.08%, 9.32%, and 9.39% of low TAWSS for ePTFE-PP, ePTFE-PB, Venous-PP, and Venous-PB anastomotic regions, respectively (**Table 5**).

**Table 5:** Haemodynamic and structural comparison of material models for the BG1 patient case. Graft and suture material vs. percentage of regions within anastomosis site (± 5 cm) with an oscillatory shear index (OSI) above 0.15, and time-averaged wall shear stresses (TAWSS) above 3 Pa and below 0.5 Pa. Mean strain and von-Mises stress of arterial model at anastomosis site (1 cm distal). Boxes highlighted yellow are above 10% of the average of their row.

| Graft / Suture Material | ePTFE / PP<br>(Stiffest) | ePTFE / PB | Vein / PP | Vein / PB<br>(Compliant) |
| --- | --- | --- | --- | --- |
| **OSI > 0.15** | 72.59% | 72.46% | 72.39% | 72.69% |
| **TAWSS > 3 Pa** | 2.70% | 2.73% | 0.49% | 0.31% |
| **TAWSS < 0.5 Pa** | 9.06% | 9.08% | 9.32% | 9.39% |
| **Mean Strain** | 4.07% | 4.19% | 5.73% | 5.97% |
| **Mean von-Mises Stress** | 284.05 kPa | 251.21 kPa | 300.33 kPa | 261.43 kPa |

In comparison with the average wall stresses of the asymptomatic patient-specific femoral-popliteal segments of 32.12 ± 4.53 kPa (AS1 to AS5), which are in good agreement with in-vivo data reported in the literature [78], average wall stresses around patient BG1’s anastomosis were elevated with 284.05 kPa, 251.21 kPa, 300.33 kPa, 261.43 kPa for ePTFE-PP, ePTFE-PB, Venous-PP, and Venous-PB anastomoses, respectively.

The use of a more compliant suture material resulted in reduced von-Mises stress of the arterial model at patient BG1’s anastomosis site (1 cm distal), while the combination of a compliant graft material with a stiff suture material resulted in the highest stresses in the arterial wall around the anastomosis (**Table 5**), which is associated with the proliferation and migration of VSMCs leading to IH and potential graft failure [79].

## 4. Discussion

Re-stenosis due to IH is one of the most common causes of mid-term vascular bypass grafts, particularly in below-the-knee applications and when for the application of synthetic graft materials. The initial trigger for the development of IH is the localised vascular trauma to the vascular wall which occurs during surgery, disrupting the endothelial and medial smooth muscle cells which in turn release various principle factors, mitogens, cytokines, and growth factors, which stimulate the quiescent vascular smooth muscle cell into a proliferative phase [80]. A range of biological, chemical, and mechanical factors have been studied with regards to their potential to aggravate the severity of IH, and compliance mismatch of graft and anastomosis with the host artery is one of the central biomechanical issues of vascular graft design, as greater compliance mismatch is associated with greater incidence and severity of IH and consequently lower mid-term patency rates.

FSI simulations are an essential tool for vascular *in silico* studies since they can be utilised to model the fluid and structural domains of the circulatory system, along with their interaction, both for native and synthetic deformable models. FSI studies are particularly useful where changes in vascular compliance, which may occur during disease or after surgical intervention, are of interest.

Biomechanical parameters which have been deemed relevant with regards to intimal hyperplasia and graft failure are wall shear stress, wall stress, and disturbed flow as characterised by high oscillatory shear index [81]. Optimising the anastomosis angle is essential to achieve favourable haemodynamic conditions. The ideal angle may vary depending on factors such as the size and type of graft, the characteristics of the native vessel, and the patient’s specific condition. *In vivo* studies have indicated that disturbed flow patterns (high OSI) and regions of excessively low or high WSS should be avoided [82,83]. An unfavourable graft-to-host-artery diameter ratio combined with a low (acute) anastomosis angle may lead to regions of high WSS just downstream of the anastomosis, which are associated with areas of rapid, unidirectional flow and may potentially lead to endothelial damage or remodelling [84]. On the other hand, obtuse angles may lead to regions of low WSS and high OSI near the anastomotic site, typically associated with areas of recirculation or stagnation, which may be a risk factor for the development of atherosclerosis and thrombosis, as it can promote endothelial dysfunction and platelet activation [85]. OSI is a measure of the directionality and magnitude of changes in shear stress during the cardiac cycle. OSI above zero are expected due to the pulsatility and partial backflow of blood flow across cardiac cycles. However, high OSI values indicate disturbed, oscillatory, or significantly reversing flow patterns, which are unphysiological in small-diameter arteries.

We observed regions of flow separation and recirculation indicated by low TAWSS and high OSI at the heel and toe of the anastomosis for all patient models, which is in good agreement with other studies and has been indicated as a risk factor for IH as significant lesions usually occur in this region [86]. While this effect may be partially attributed to the geometric composition of end-to-side anastomoses, low-WSS recirculation zones have also been reported in idealised end-to-end FSI studies where blood flows from a low-compliance into a high-compliance vessel, i.e., at the distal anastomosis between a host artery and stiff graft or proximal anastomosis between a host artery and a graft with a mechanical compliance that exceeds arterial compliance [87].

Our numerical results demonstrate non-physiological flow patterns in the anastomotic region. On the artery floor opposite the junction flow separation and zones of recirculation were found. Fluctuating WSS as indicated by a high OSI can increase the incidence and severity of IH through multiple signalling pathways, in which oxidative stress factors are implicated, promoting the proliferation, migration, and increased survival of VSMCs in the arterial intima [88]. High OSI is associated with rapidly changing flow directions and high spatial and temporal WSS gradients, which may cause injury and exfoliation of the endothelium, thus triggering the proliferation and migration of VSMCs and causing IH [88,89].

Furthermore, our results indicate an increase in graft compliance results in a decrease in adversely high TAWSS (TAWSS > 3 Pa) and a slight increase in adversely low TAWSS (< 0.5 Pa). This can be expected, as a larger vessel diameter – e.g., linked to an increased compliance – is likely to result in a lower WSS, if the cardiac output is constant. FSI models of vascular anastomoses comparing deformable and rigid vascular wall have shown that the deformation of the arterial wall has a damping effect which reduces peak wall shear stresses, but the anastomotic compliance mismatch between the arterial and graft model induces more oscillatory WSS which may contribute to the initiation of intimal hyperplasia, indicating the presence of a stiff graft within the circulatory system counteracts the protective function of the compliant arterial wall [90]. However, the results of this study are limited by the lack of patient-specific boundary conditions since no clinical velocity and pressure data were available for this post-hoc analysis. Furthermore, the arterial wall and graft wall were assumed to be single-layer structures with uniform thicknesses since detailed wall models could not be extracted from the medical imaging data sets.

Our results indicate that for both PP and saphenous vein suture materials, the arterial wall movement is restricted compared with a pristine vessel, resulting in elevated stresses at the arterial wall close to the anastomosis. While the effect of the suture material on haemodynamic parameters such as WSS and OSI is minimal, the suture material compliance affects the local von-Mises stresses in the arterial wall. While the use of the more compliant PB suture material resulted in reduced von-Mises stress of the arterial model at the anastomosis site (1 cm distal), the combination of a compliant graft material with a stiff suture material resulted in the highest stresses in the arterial wall at the anastomosis. Since stress concentrations occur in connections between materials of different stiffnesses, a stiff suture material connected to two compliant graft materials leads to a stricture effect at the anastomosis with stress peaks. Conversely, optimising anastomotic compliance appears to be particularly relevant in combination with compliant grafts to minimise adverse arterial wall stresses. When compared with published *in vivo* animal studies on intimal hyperplasia, correlations emerge between suture line stresses and anastomotic intimal hyperplasia, with a pronounced compliance mismatch appearing to exert a proliferative effect on suture line hyperplasia [91].

## 5. Conclusions

This study describes a framework to simulate complex interactions between peripheral blood flow, femoropopliteal artery walls, and/or grafted material structures which are reconstructed from CTA imagery of five patients who have had a side-to-end bypass graft and five healthy patients. Non-Newtonian fluid model and three structural models were applied to simulate the artery, the graft, and the suture material. Furthermore, all geometries were pre-stressed to ensure our FSI models were physiologically pressurised when simulations begun. Our patient database included graft-to-artery anastomosis angles varying between 21**°** and 36**°**, where our simulations predict bypass graft patients contained larger areas of low vessel wall shear stress (p = 0.005) and increasing the bypass anastomotic angle produces larger regions of oscillatory blood flow (R^2^ = 0.96). Our multi-structural FSI simulation allowed us to evaluate the effect of different implant and suture materials on blood flow, where we discovered that compliant graft and suture materials are both required to reduce nonphysiologically high shear stresses downstream of the anastomosis site. Altogether, minimizing anastomosis angle and using mechanically-compliant grafts and sutures together will reduce regions of nonphysiologically low wall shear stress and high wall stresses, which are known risk factors leading to peripheral bypass intimal hyperplasia and restenosis. Our study provides a framework to simulate patient-specific fluid-structure interactions for peripheral artery grafts and in doing so provides a valuable tool for preoperative surgical planning.

## Statement of Ethical Approval

Non-identifiable patient images were provided under human research ethics approval LNR/2019/QRBW/53686 and site-specific approval SSA/QMS/53686 “Vascular model biofabrication for surgical planning, training, and treatment.”

## Supplemental Information

**Table S1:** Fourier coefficients for pulsatile velocity inlet and pressure outlet.

| Parameter | Velocity Profile (95% confidence) | Pressure Profile (95% confidence) |
| --- | --- | --- |
| $a_0$ | 0.1731 (0.1728, 0.1734) | $1.267 \times 10^4$ ($1.267 \times 10^4$, $1.267 \times 10^4$) |
| $a_1$ | 0.1129 (0.1124, 0.1133) | -330.4 (-335.1, -325.8) |
| $b_1$ | 0.122 (0.1216, 0.1224) | 2877 (2873, 2881) |
| $a_2$ | -0.02976 (-0.03032, -0.0292) | -1196 (-1201, -1192) |
| $b_2$ | 0.1468 (0.1464, 0.1472) | 987.6 (983.1, 992.1) |
| $a_3$ | -0.05254 (-0.05295, -0.05212) | -524.8 (-528.4, -521.1) |
| $b_3$ | 0.03393 (0.0335, 0.03436) | 68 (64.11, 71.9) |
| $a_4$ | -0.03287 (-0.03328, -0.03246) | -309 (-312.7, -305.3) |
| $b_4$ | 0.0153 (0.01488, 0.01571) | 78.97 (75.16, 82.79) |
| $a_5$ | -0.02275 (-0.02315, -0.02235) | -222.6 (-226.3, -218.9) |
| $b_5$ | -0.006089 (-0.006502, -0.005677) | -82.71 (-86.52, -78.9) |
| $a_6$ | -0.004072 (-0.004471, -0.003673) | -47.45 (-51.13, -43.76) |
| $b_6$ | -0.003696 (-0.004095, -0.003296) | -53.38 (-57.07, -49.69) |
| $a_7$ | -0.003927 (-0.004327, -0.003527) | -45.23 (-48.92, -41.54) |
| $b_7$ | -0.0003096 (-0.000709, $8.979 \times 10^{-5}$) | -13.24 (-16.93, -9.542) |
| $a_8$ | -0.001664 (-0.002063, -0.001265) | -19 (-22.69, -15.32) |
| $b_8$ | -0.001678 (-0.002077, -0.001278) | -28.65 (-32.34, -24.96) |
| $\omega$ | 7.994 (7.992, 7.996) | 7.994 (7.992, 7.996) |

**Table S2:** Haemodynamic comparison of ePTFE-PP patient cases. Anastomosis angle vs. percentage of regions within anastomosis site (± 5 cm) with an oscillatory shear index (OSI) above 0.15, and time-averaged wall shear stresses (TAWSS) above 3 Pa and below 0.5 Pa.

| Patient Case | BG1 | BG2 | BG3 | BG4 | BG5 |
| --- | --- | --- | --- | --- | --- |
| **Anastomosis Angle** | 35.4° | 28.9° | 28.3° | 21.1° | 32.2° |
| **OSI > 0.15** | 72.59% | 43.49% | 49.19% | 25.29% | 60.24% |
| **TAWSS > 3 Pa** | 2.70% | 0.59% | 0.00% | 13.74% | 0.00% |
| **TAWSS < 0.5 Pa** | 9.06% | 7.55% | 13.42% | 8.13% | 14.62% |

**Table S3:** Haemodynamic and structural summary of asymptomatic patient cases. Time-averaged wall stresses (TAWS), percentage of regions with an oscillatory shear index (OSI) above 0.15, and time-averaged wall shear stresses (TAWSS) above 3 Pa and below 0.5 Pa.

| Patient Case | AS1 | AS2 | AS3 | AS4 | AS5 |
| --- | --- | --- | --- | --- | --- |
| **TAWSS [kPa]** | 39.38 | 28.71 | 27.91 | 29.11 | 35.50 |
| **OSI > 0.15** | 18.01% | 7.69% | 23.85% | 2.49% | 7.57% |
| **TAWSS > 3 Pa** | 0.00% | 0.55% | 5.22% | 4.10% | 2.87% |
| **TAWSS < 0.5 Pa** | 0.00% | 0.00% | 0.00% | 0.00% | 0.00% |

### Mesh Convergence

#### Bypass Graft – BG1

Structural
|  |  |  |
| --- | --- | --- |
| **Point_Suture_Min** |  |  |
| **Conduit_Anastomosis Element** | **Deformation [m]** | **von-Mises Stress [Pa]** |
| 0.0007_0.00035 | 1.08225E-05 | 24686.746 |
| Change [%] | −0.95 | 42.19 |
| 0.00055_0.00025 | 1.07205E-05 | 42706.569 |
| Change [%] | 0.23 | −8.56 |
| 0.00045_0.0002 | 1.06962E-05 | 46362.355 |
| Change [%] | 0.034 | 1.65 |
| 0.0004_0.00015 | 1.06999E-05 | 47139.021 |
| **Point_Suture_Max** |  |  |
| **Conduit_Anastomosis Element** | **Deformation [m]** | **von-Mises Stress [Pa]** |
| 0.0007_0.00035 | 8.43552E-06 | 145659.56 |
| Change [%] | 0.22 | 14.282 |
| 0.00055_0.00025 | 8.45397E-06 | 169933.41 |
| Change [%] | −0.38 | 0.66 |
| 0.00045_0.0002 | 8.48642E-06 | 168824 |
| Change [%] | 0.49 | 1.27 |
| 0.0004_0.00015 | 8.52789E-06 | 170993.33 |
| **Point_Graft** |  |  |
| **Conduit_Anastomosis Element** | **Deformation [m]** | **von-Mises Stress [Pa]** |
| 0.0007_0.00035 | 5.19978E-06 | 3560.7754 |
| Change [%] | −1.86 | 5.63 |
| 0.00055_0.00025 | 5.10506E-06 | 3773.1939 |
| Change [%] | 0.56 | 2.82 |
| 0.00045_0.0002 | 5.07623E-06 | 3882.4932 |
| Change [%] | 0.40 | −0.57 |
| 0.0004_0.00015 | 5.0968E-06 | 3860.5991 |
| **Point_Artery** |  |  |
| **Conduit_Anastomosis Element** | **Deformation [m]** | **von-Mises Stress [Pa]** |
| 0.0007_0.00035 | 1.49308E-05 | 1799.684 |
| Change [%] | 1.08 | −1.65 |
| 0.00055_0.00025 | 1.5094E-05 | 1770.4654 |
| Change [%] | −0.028 | 0.967 |
| 0.00045_0.0002 | 1.50982E-05 | 1753.3572 |
| Change [%] | 1.25 | 1.56 |
| 0.0004_0.00015 | 1.52889E-05 | 1781.1801 |

*Fluid*
| # Nodes | Velocity [m/s] | Change [%] | Pressure [Pa] | Change [%] | Wall Shear Stress [Pa] | Change [%] |
| --- | --- | --- | --- | --- | --- | --- |
| 385k | 0.1888 |  | 12652.8066 |  | 0.1017 |  |
|  |  | 2.10762 |  | 0.022 |  | -6.64 |
| 579k | 0.1929 |  | 12655.5215 |  | 0.0954 |  |
|  |  | 0.83424 |  | -0.014 |  | 5.93 |
| 901k | 0.1945 |  | 12653.7568 |  | 0.1014 |  |
|  |  | 0.82358 |  | -0.0007 |  | 0.59 |
| 1,372k | 0.1961 |  | 12653.6650 |  | 0.1020 |  |
|  |  | 0.0928 |  | -0.0055 |  | 1.57 |
| 2,315k | 0.1963 |  | 12652.9668 |  | 0.1036 |  |

#### Bypass Graft – BG2

**Bypass Graft – BG2***Structural*
|  |  |  |
| --- | --- | --- |
| **Point_Suture_Min** |  |  |
| **Conduit_Anastomosis Element** | **Deformation [m]** | **von-Mises Stress [Pa]** |
| 0.0007_0.00035 | 2.08481E-05 | 25936.99 |
| Change [%] | 0.22 | -15.90 |
| 0.00055_0.00025 | 2.08935E-05 | 22378.02 |
| Change [%] | 0.030 | 0.33 |
| 0.00045_0.0002 | 2.08872E-05 | 22304.41 |
| Change [%] | 0.089 | 1.09 |
| 0.0004_0.00015 | 2.09057E-05 | 22549.42 |
| **Point_Suture_Max** |  |  |
| **Conduit_Anastomosis Element** | **Deformation [m]** | **von-Mises Stress [Pa]** |
| 0.0007_0.00035 | 2.16564E-05 | 180094.3 |
| Change [%] | 0.99 | 10.89 |
| 0.00055_0.00025 | 2.18733E-05 | 202103.7 |
| Change [%] | -1.27 | -9.59 |
| 0.00045_0.0002 | 2.21520E-05 | 221486.1 |
| Change [%] | 0.77 | -0.76 |
| 0.0004_0.00015 | 2.23230E-05 | 219814.3 |
| **Point_Graft** |  |  |
| **Conduit_Anastomosis Element** | **Deformation [m]** | **von-Mises Stress [Pa]** |
| 0.0007_0.00035 | 2.90944E-06 | 3948.871 |
| Change [%] | -2.52 | -2.10 |
| 0.00055_0.00025 | 2.83783E-06 | 3867.77 |
| Change [%] | 0.42306984 | -1.06083 |
| 0.00045_0.0002 | 2.82582E-06 | 3827.17 |
| Change [%] | −0.42 | 2.18 |
| 0.0004_0.00015 | 2.81402E-06 | 3912.327 |
| **Point_Artery** |  |  |
| **Conduit_Anastomosis Element** | **Deformation [m]** | **von-Mises Stress [Pa]** |
| 0.0007_0.00035 | 1.54254E-05 | 1839.058 |
| Change [%] | 0.64 | −2.25 |
| 0.00055_0.00025 | 1.55251E-05 | 1798.621 |
| Change [%] | −1.00 | 0.98 |
| 0.00045_0.0002 | 1.56806E-05 | 1781.071 |
| Change [%] | 0.80 | 0.60 |
| 0.0004_0.00015 | 1.58065E-05 | 1791.896 |

*Fluid*
| # Nodes | Velocity [m/s] | Change [%] | Pressure [Pa] | Change [%] | Wall Shear Stress [Pa] | Change [%] |
| --- | --- | --- | --- | --- | --- | --- |
| 411k | 0.3481 |  | 12735.9375 |  | 1.0739 |  |
|  |  | 0.018 |  | −0.0076 |  | 2.47 |
| 676k | 0.3482 |  | 12734.9648 |  | 1.1012 |  |
|  |  | 0.27 |  | −0.0006 |  | 1.52 |
| 1055k | 0.3472 |  | 12734.8867 |  | 1.1182 |  |
|  |  | 0.024 |  | 0.0077 |  | 1.31 |
| 1603k | 0.3472 |  | 12735.8643 |  | 1.1331 |  |
|  |  | 0.74 |  | −0.0079 |  | −0.09 |
| 2796k | 0.3446 |  | 12734.8535 |  | 1.1321 |  |

#### Bypass Graft – BG3

Structural
|  |  |  |
| --- | --- | --- |
| **Point_Suture_Min** |  |  |
| **Conduit_Anastomosis Element** | **Deformation [m]** | **von-Mises Stress [Pa]** |
| 0.0007_0.00035 | 1.69633E-05 | 64501.53 |
| Change [%] | 0.31 | –5.99 |
| 0.00055_0.00025 | 1.70157E-05 | 60854.78 |
| Change [%] | –0.64 | 0.16 |
| 0.00045_0.0002 | 1.71249E-05 | 60757.82 |
| Change [%] | 0.48 | 1.14 |
| 0.0004_0.00015 | 1.72072E-05 | 61459.02 |
| **Point_Suture_Max** |  |  |
| **Conduit_Anastomosis Element** | **Deformation [m]** | **von-Mises Stress [Pa]** |
| 0.0007_0.00035 | 2.29169E-05 | 155963.8 |
| Change [%] | 0.88 | 0.84 |
| 0.00055_0.00025 | 2.31215E-05 | 157288 |
| Change [%] | –0.41 | 0.71 |
| 0.00045_0.0002 | 2.32153E-05 | 156167.6 |
| Change [%] | 0.36 | 1.17 |
| 0.0004_0.00015 | 2.32986E-05 | 158010.9 |
| **Point_Graft** |  |  |
| **Conduit_Anastomosis Element** | **Deformation [m]** | **von-Mises Stress [Pa]** |
| 0.0007_0.00035 | 4.12231E-06 | 5045.176 |
| Change [%] | –1.00 | 1.59 |
| 0.00055_0.00025 | 4.08148E-06 | 5126.627 |
| Change [%] | 0.46 | 0.26 |
| 0.00045_0.0002 | 4.06288E-06 | 5139.888 |
| Change [%] | –0.24 | –0.60 |
| 0.0004_0.00015 | 4.05315E-06 | 5109.43 |
| **Point_Artery** |  |  |
| **Conduit_Anastomosis Element** | **Deformation [m]** | **von-Mises Stress [Pa]** |
| 0.0007_0.00035 | 1.68524E-05 | 1855.736 |
| Change [%] | 0.37 | 3.06 |
| 0.00055_0.00025 | 1.69152E-05 | 1914.24 |
| Change [%] | –0.066 | –0.35 |
| 0.00045_0.0002 | 1.69263E-05 | 1921.019 |
| Change [%] | 0.51 | –0.32 |
| 0.0004_0.00015 | 1.70124E-05 | 1914.916 |

*Fluid*
| # Nodes | Velocity [m/s] | Change [%] | Pressure [Pa] | Change [%] | Wall Shear Stress [Pa] | Change [%] |
| --- | --- | --- | --- | --- | --- | --- |
| 504k | 0.2085 |  | 12672.0781 |  | 1.5519 |  |
|  |  | 6.14 |  | -0.0069 |  | 0.33 |
| 770k | 0.2222 |  | 12671.2090 |  | 1.5570 |  |
|  |  | 2.34 |  | -0.0091 |  | -1.93 |
| 1,172k | 0.2275 |  | 12670.0498 |  | 1.5276 |  |
|  |  | 1.58 |  | 0.0031 |  | 1.72 |
| 1,862k | 0.2311 |  | 12670.4434 |  | 1.5544 |  |
|  |  | 1.15 |  | -0.0044 |  | -0.017 |
| 2,992k | 0.2338 |  | 12669.8857 |  | 1.5541 |  |

#### Bypass Graft – BG4

**Bypass Graft – BG4***Structural*
|  |  |  |
| --- | --- | --- |
| **Point_Suture_Min** |  |  |
| **Conduit_Anastomosis Element** | **Deformation [m]** | **von-Mises Stress [Pa]** |
| 0.0007_0.00035 | 6.00293E-06 | 97378.88 |
| Change [%] | 1.44 | 32.72 |
| 0.00055_0.00025 | 6.09094E-06 | 144747.4 |
| Change [%] | -1.84 | 3.41 |
| 0.00045_0.0002 | 6.20293E-06 | 139806.9 |
| Change [%] | 0.24 | 0.58 |
| 0.0004_0.00015 | 6.21771E-06 | 140622.7 |
| **Point_Suture_Max** |  |  |
| **Conduit_Anastomosis Element** | **Deformation [m]** | **von-Mises Stress [Pa]** |
| 0.0007_0.00035 | 6.00293E-06 | 114516.6 |
| Change [%] | 12.73 | 31.96 |
| 0.00055_0.00025 | 6.87845E-06 | 168316.8 |
| Change [%] | -1.33 | 0.50 |
| 0.00045_0.0002 | 6.96984E-06 | 167469 |
| Change [%] | 0.31 | -0.38 |
| 0.0004_0.00015 | 6.99172E-06 | 166835.9 |
| **Point_Graft** |  |  |
| **Conduit_Anastomosis Element** | **Deformation [m]** | **von-Mises Stress [Pa]** |
| 0.0007_0.00035 | 9.40356E-07 | 3290.756 |
| Change [%] | 5.83 | -10.42 |
| 0.00055_0.00025 | 9.98618E-07 | 2980.117 |
| Change [%] | 2.26 | 1.73 |
| 0.00045_0.0002 | 9.76000E-07 | 3032.662 |
| Change [%] | -0.42 | -1.89 |
| 0.0004_0.00015 | 9.71932E-07 | 2976.32 |
| **Point_Artery** |  |  |
| **Conduit_Anastomosis Element** | **Deformation [m]** | **von-Mises Stress [Pa]** |
| 0.0007_0.00035 | 1.05751E-05 | 1560.057 |
| Change [%] | 2.95 | -1.22 |
| 0.00055_0.00025 | 1.08963E-05 | 1541.295 |
| Change [%] | -0.060 | 1.60 |
| 0.00045_0.0002 | 1.09028E-05 | 1516.657 |
| Change [%] | -0.62 | -1.52 |
| 0.0004_0.00015 | 1.08357E-05 | 1494.022 |

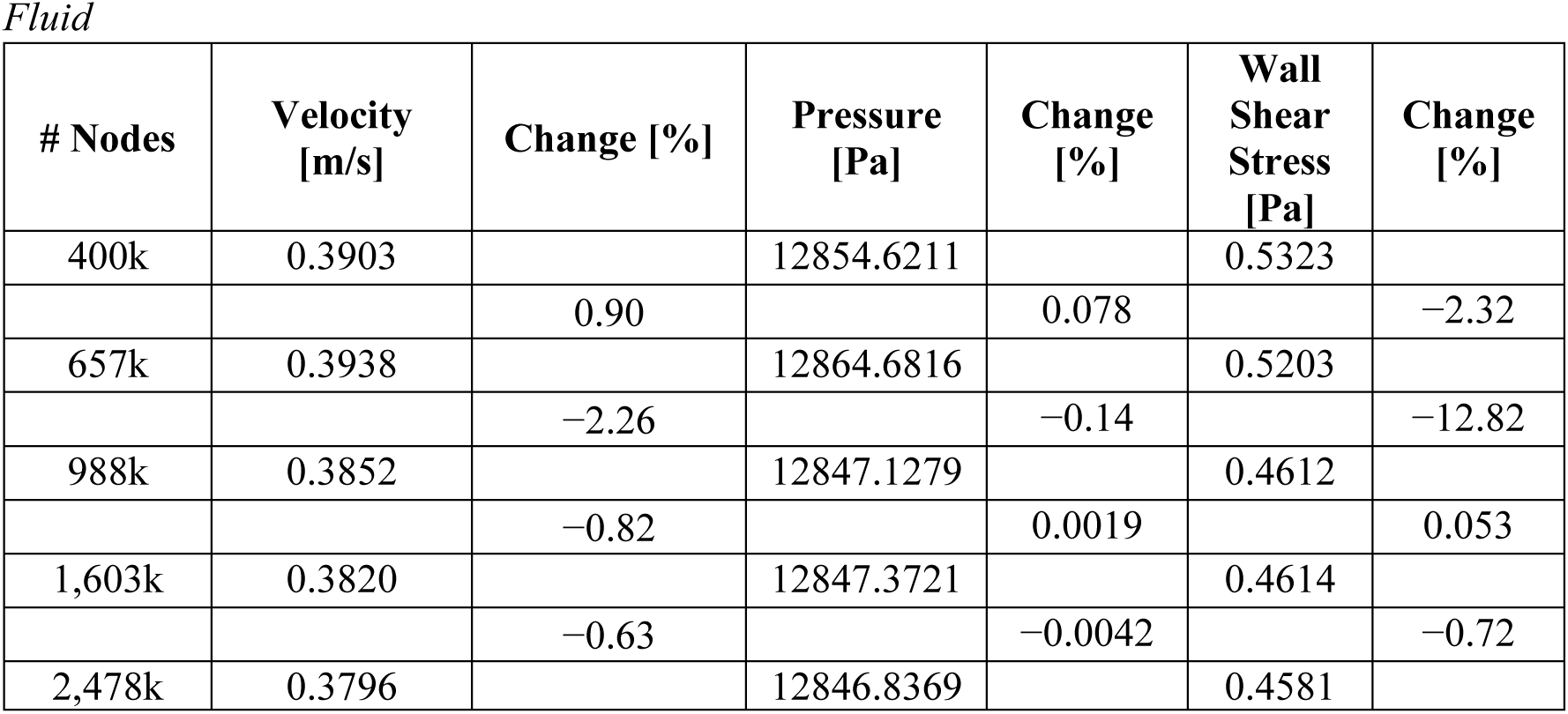

#### Bypass Graft – BG5

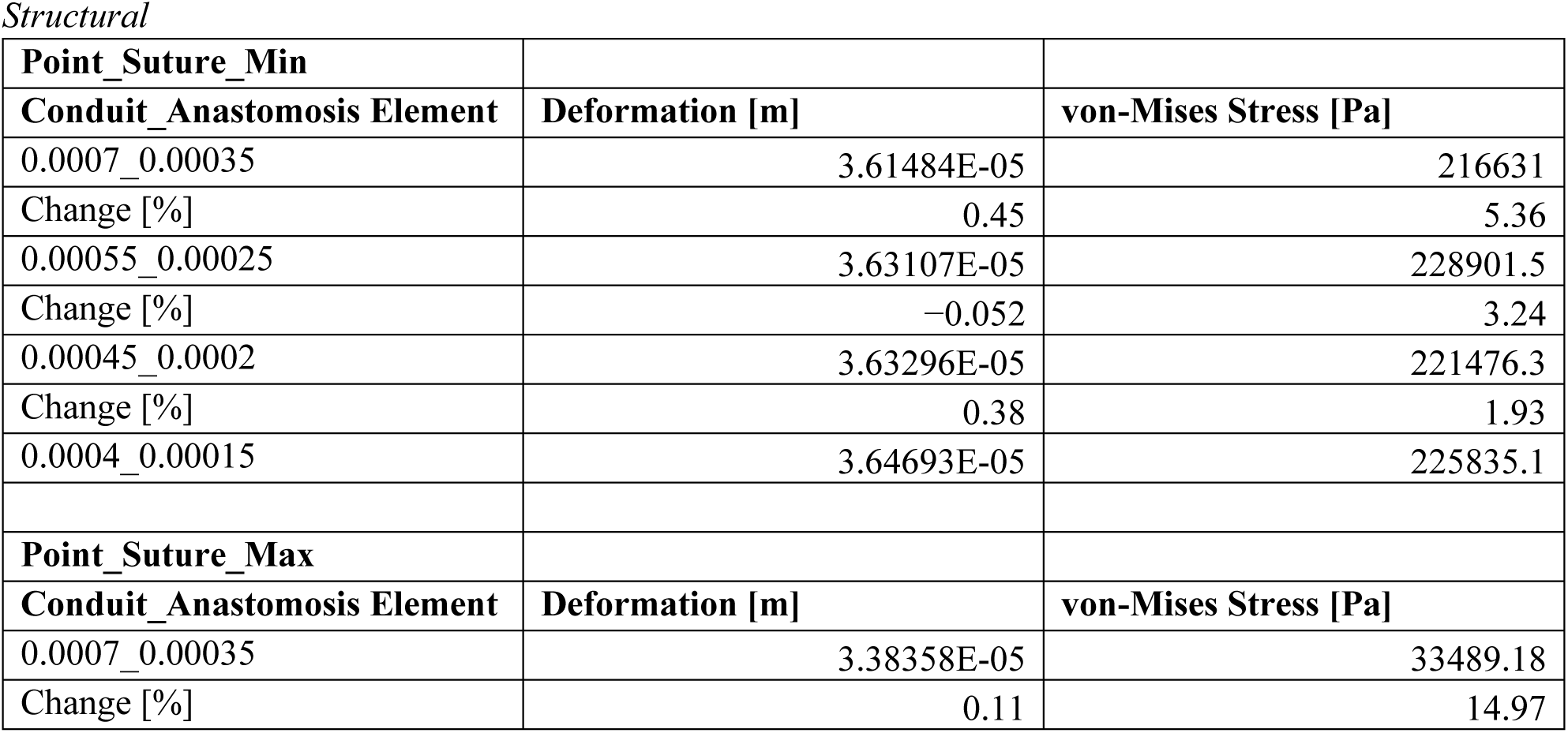
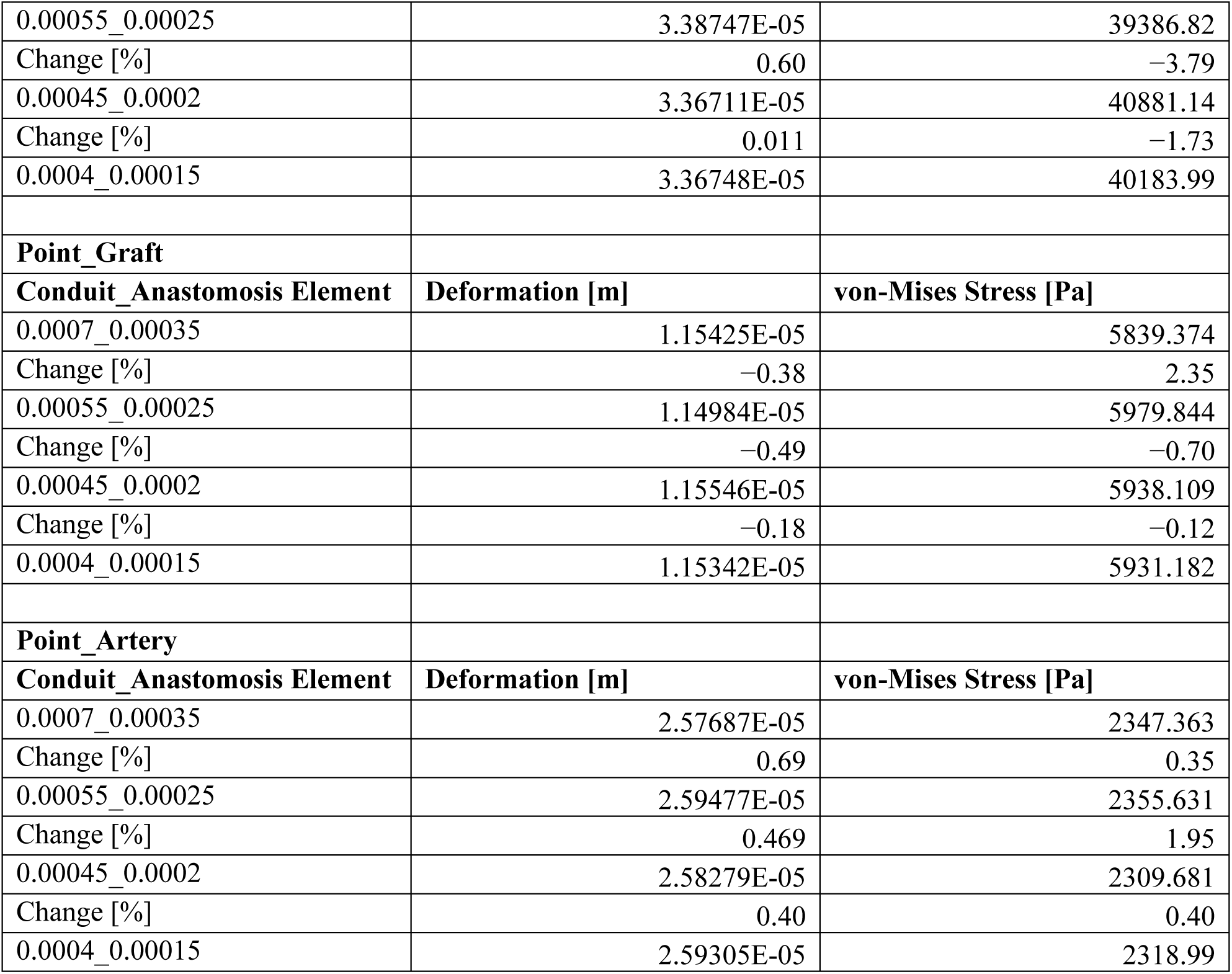

*Fluid*
| # Nodes | Velocity [m/s] | Change [%] | Pressure [Pa] | Change [%] | Wall Shear Stress [Pa] | Change [%] |
| --- | --- | --- | --- | --- | --- | --- |
| 391k | 0.0150 |  | 12662.5342 |  | 0.0751 |  |
|  |  | -1.24 |  | -0.0070 |  | -3.55 |
| 606k | 0.0148 |  | 12661.6504 |  | 0.0725 |  |
|  |  | -1.13 |  | -0.0066 |  | -1.45 |
| 955k | 0.0147 |  | 12660.8105 |  | 0.0715 |  |
|  |  | 0.33 |  | -0.0042 |  | -1.20 |
| 1,485k | 0.0147 |  | 12660.2754 |  | 0.0706 |  |
|  |  | -0.92 |  | 0.0049 |  | -0.72 |
| 2,301k | 0.0146 |  | 12660.8965 |  | 0.0701 |  |

### Asymptomatic – AS1

Asymptomatic – AS1 Structural
| # Nodes | Deformation [m] | Change [%] | von-Mises Stress [Pa] | Change [%] |
| --- | --- | --- | --- | --- |
| 131k | 1.73530E-05 |  | 1888.4 |  |
|  |  | −0.17 |  | 1.43 |
| 208k | 1.73240E-05 |  | 1915.7 |  |
|  |  | 0.74 |  | −3.54 |
| 316k | 1.74530E-05 |  | 1850.2 |  |
|  |  | 0.22 |  | −0.25 |
| 480k | 1.74920E-05 |  | 1845.6 |  |
|  |  | −0.70 |  | 1.17 |
| 725k | 1.73700E-05 |  | 1867.5 |  |

Fluid
| # Nodes | Velocity [m/s] | Change [%] | Pressure [Pa] | Change [%] | Wall Shear Stress [Pa] | Change [%] |
| --- | --- | --- | --- | --- | --- | --- |
| 140k | 0.3113 |  | 12706.0742 |  | 0.9837 |  |
|  |  | 1.27 |  | 0.0025 |  | −2.20 |
| 212k | 0.3074 |  | 12706.3896 |  | 0.9626 |  |
|  |  | 0.59 |  | 0.0025 |  | 1.16 |
| 318k | 0.3093 |  | 12706.7051 |  | 0.9739 |  |
|  |  | −0.26 |  | −0.0003 |  | −1.07 |
| 480k | 0.3085 |  | 12706.6719 |  | 0.9636 |  |
|  |  | −0.33 |  | −0.0022 |  | −0.11 |
| 728k | 0.3074 |  | 12706.3896 |  | 0.9626 |  |

### Asymptomatic – AS2

Asymptomatic – AS2 Structural
| # Nodes | Deformation [m] | Change [%] | von-Mises Stress [Pa] | Change [%] |
| --- | --- | --- | --- | --- |
| 131k | 8.77148E-06 |  | 1201.901 |  |
|  |  | 2.39 |  | 7.98 |
| 203k | 8.98671E-06 |  | 1306.1641 |  |
|  |  | −0.44 |  | −4.14 |
| 311k | 8.94770E-06 |  | 1254.1796 |  |
|  |  | 0.21 |  | 1.51 |
| 469k | 8.96665E-06 |  | 1273.3572 |  |
|  |  | −0.054 |  | −1.22 |
| 704k | 8.96184E-06 |  | 1258.0683 |  |

Fluid
| # Nodes | Velocity [m/s] | Change [%] | Pressure [Pa] | Change [%] | Wall Shear Stress [Pa] | Change [%] |
| --- | --- | --- | --- | --- | --- | --- |
| 145k | 0.4103 |  | 12764.8271 |  | 1.3961 |  |
|  |  | 1.01 |  | 0.031 |  | −6.14 |
| 218k | 0.4144 |  | 12768.7578 |  | 1.3154 |  |
|  |  | 1.09 |  | 0.0011 |  | 0.42 |
| 335k | 0.4190 |  | 12768.8975 |  | 1.3210 |  |
|  |  | 1.11 |  | 0.0004 |  | −0.55 |
| 502k | 0.4237 |  | 12768.9434 |  | 1.3137 |  |
|  |  | 0.48 |  | −0.0037 |  | −1.51 |
| 771k | 0.4257 |  | 12768.4707 |  | 1.2942 |  |

### Asymptomatic – AS3

Asymptomatic – AS3 Structural
| # Nodes | Deformation [m] | Change [%] | von-Mises Stress [Pa] | Change [%] |
| --- | --- | --- | --- | --- |
| 133k | 1.09417E-05 |  | 1357.2851 |  |
|  |  | 1.19 |  | –3.50 |
| 205k | 1.10729E-05 |  | 1311.4256 |  |
|  |  | –0.26 |  | 1.59 |
| 309k | 1.10443E-05 |  | 1332.6343 |  |
|  |  | –0.098 |  | 0.77 |
| 485k | 1.10335E-05 |  | 1342.9247 |  |
|  |  | 0.54 |  | –1.28 |
| 730k | 1.10934E-05 |  | 1325.891 |  |

Fluid
| # Nodes | Velocity [m/s] | Change [%] | Pressure [Pa] | Change [%] | Wall Shear Stress [Pa] | Change [%] |
| --- | --- | --- | --- | --- | --- | --- |
| 131k | 0.3761 |  | 12828.1211 |  | 1.2175 |  |
|  |  | 2.39 |  | 0.0043 |  | 0.22 |
| 201k | 0.3853 |  | 12828.6699 |  | 1.2201 |  |
|  |  | 1.28 |  | 0.0024 |  | –2.08 |
| 310k | 0.3903 |  | 12828.9785 |  | 1.1952 |  |
|  |  | 0.61 |  | –0.0002 |  | –0.93 |
| 468k | 0.3927 |  | 12828.9531 |  | 1.1842 |  |
|  |  | 0.19 |  | 0.0013 |  | 1.06 |
| 712k | 0.3935 |  | 12829.1182 |  | 1.1969 |  |

### Asymptomatic – AS4

Asymptomatic – AS4 Structural
| # Nodes | Deformation [m] | Change [%] | von-Mises Stress [Pa] | Change [%] |
| --- | --- | --- | --- | --- |
| 136k | 1.95810E-05 |  | 1974.8 |  |
|  |  | 1.08 |  | 0.70 |
| 206k | 1.97940E-05 |  | 1988.8 |  |
|  |  | 0.72 |  | 0.42 |
| 313k | 1.99380E-05 |  | 1997.1 |  |
|  |  | −0.015 |  | −0.71 |
| 471k | 1.99350E-05 |  | 1983.0 |  |
|  |  | 0.035 |  | −0.61 |
| 713k | 1.99420E-05 |  | 1970.9 |  |

Fluid
| # Nodes | Velocity [m/s] | Change [%] | Pressure [Pa] | Change [%] | Wall Shear Stress [Pa] | Change [%] |
| --- | --- | --- | --- | --- | --- | --- |
| 132k | 0.3140 |  | 12721.3184 |  | 1.4663 |  |
|  |  | 2.69 |  | 0.0066 |  | −0.055 |
| 201k | 0.3227 |  | 12722.1543 |  | 1.4655 |  |
|  |  | 1.115 |  | 0.0006 |  | −1.01 |
| 304k | 0.3263 |  | 12722.2305 |  | 1.4509 |  |
|  |  | 0.078 |  | −0.0001 |  | 0.95 |
| 458k | 0.3266 |  | 12722.2188 |  | 1.4648 |  |
|  |  | −0.039 |  | −0.0002 |  | −1.15 |
| 691k | 0.3265 |  | 12722.1953 |  | 1.4481 |  |

### Asymptomatic – AS5

Asymptomatic – AS5 Structural
| # Nodes | Deformation [m] | Change [%] | von-Mises Stress [Pa] | Change [%] |
| --- | --- | --- | --- | --- |
| 131k | 1.10030E-05 |  | 2646.8 |  |
|  |  | 0.27 |  | −3.67 |
| 208k | 1.10330E-05 |  | 2553.1 |  |
|  |  | 0.027 |  | 2.07 |
| 316k | 1.10360E-05 |  | 2607 |  |
|  |  | −0.23 |  | 0.015 |
| 480k | 1.10110E-05 |  | 2607.4 |  |
|  |  | −0.40 |  | 0.67 |
| 725k | 1.09670E-05 |  | 2624.9 |  |

Fluid
| # Nodes | Velocity [m/s] | Change [%] | Pressure [Pa] | Change [%] | Wall Shear Stress [Pa] | Change [%] |
| --- | --- | --- | --- | --- | --- | --- |
| 126k | 0.3839 |  | 12735.9902 |  | 1.7386 |  |
|  |  | 3.01 |  | 0.0085 |  | −5.44 |
| 196k | 0.3958 |  | 12737.0791 |  | 1.6489 |  |
|  |  | 0.66 |  | −0.013 |  | 3.43 |
| 297k | 0.3985 |  | 12735.3799 |  | 1.7076 |  |
|  |  | 0.98 |  | 0.0006 |  | −0.62 |
| 448k | 0.4024 |  | 12735.4541 |  | 1.6970 |  |
|  |  | 0.42 |  | −0.0011 |  | 0.71 |
| 681k | 0.4041 |  | 12735.3145 |  | 1.7091 |  |

### Timestep Convergence

#### Velocity

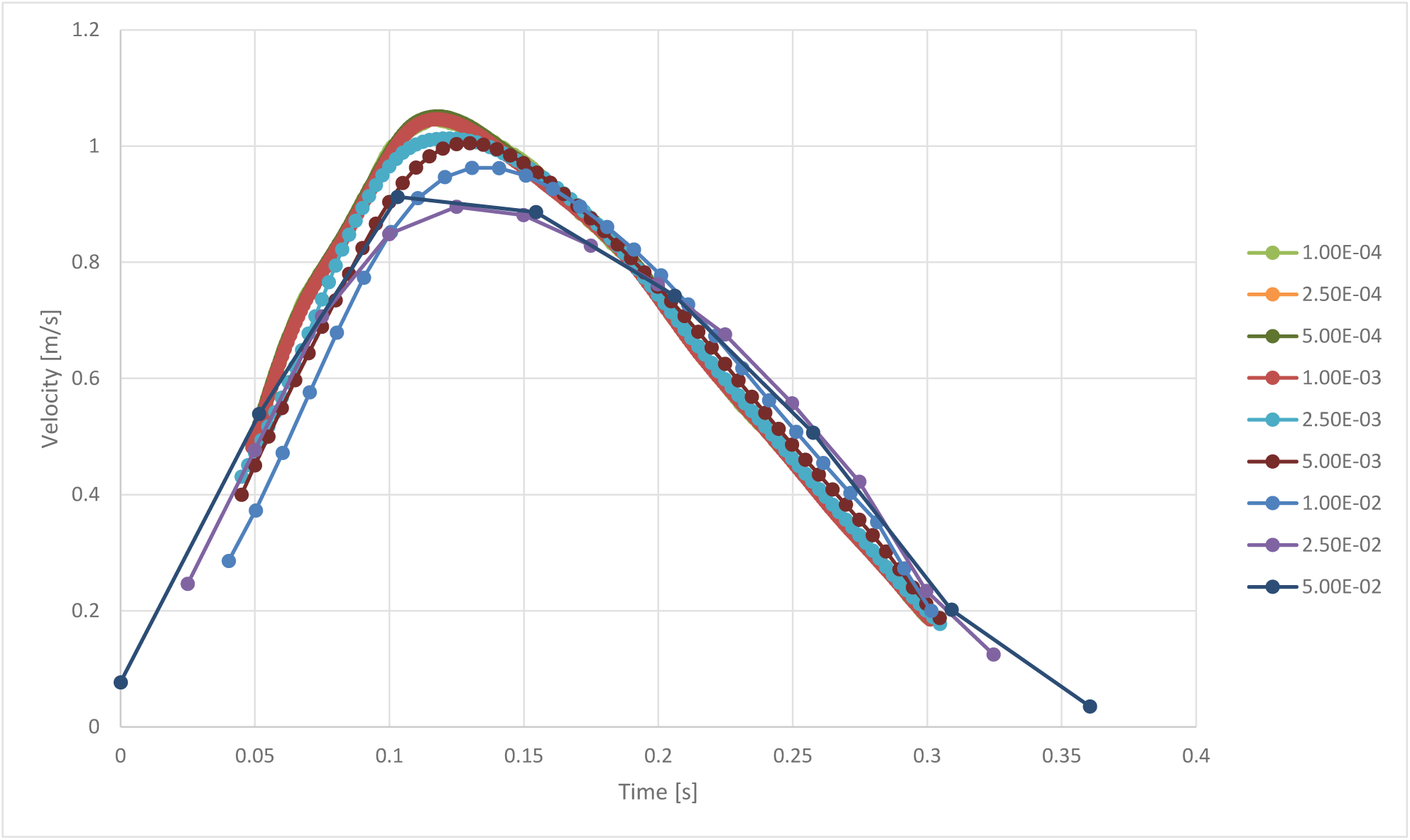

| Timepoint: 0.12 s |  |  |
| --- | --- | --- |
| Timestep [s] | Velocity [m/s] | Relative Change [%] |
| 5.00E-02 | 0.904058 |  |
|  |  | 1.98 |
| 2.50E-02 | 0.886492 |  |
|  |  | −6.15 |
| 1.00E-02 | 0.944583 |  |
|  |  | −5.18 |
| 5.00E-03 | 0.99615 |  |
|  |  | −1.68 |
| 2.50E-03 | 1.01318 |  |
|  |  | −3.09 |
| 1.00E-03 | 1.045498 |  |
|  |  | −0.50 |
| 5.00E-04 | 1.05072 |  |
|  |  | 0.13 |
| 2.50E-04 | 1.049317 |  |
|  |  | 0.53 |
| 1.00E-04 | 1.043779 |  |

#### Pressure

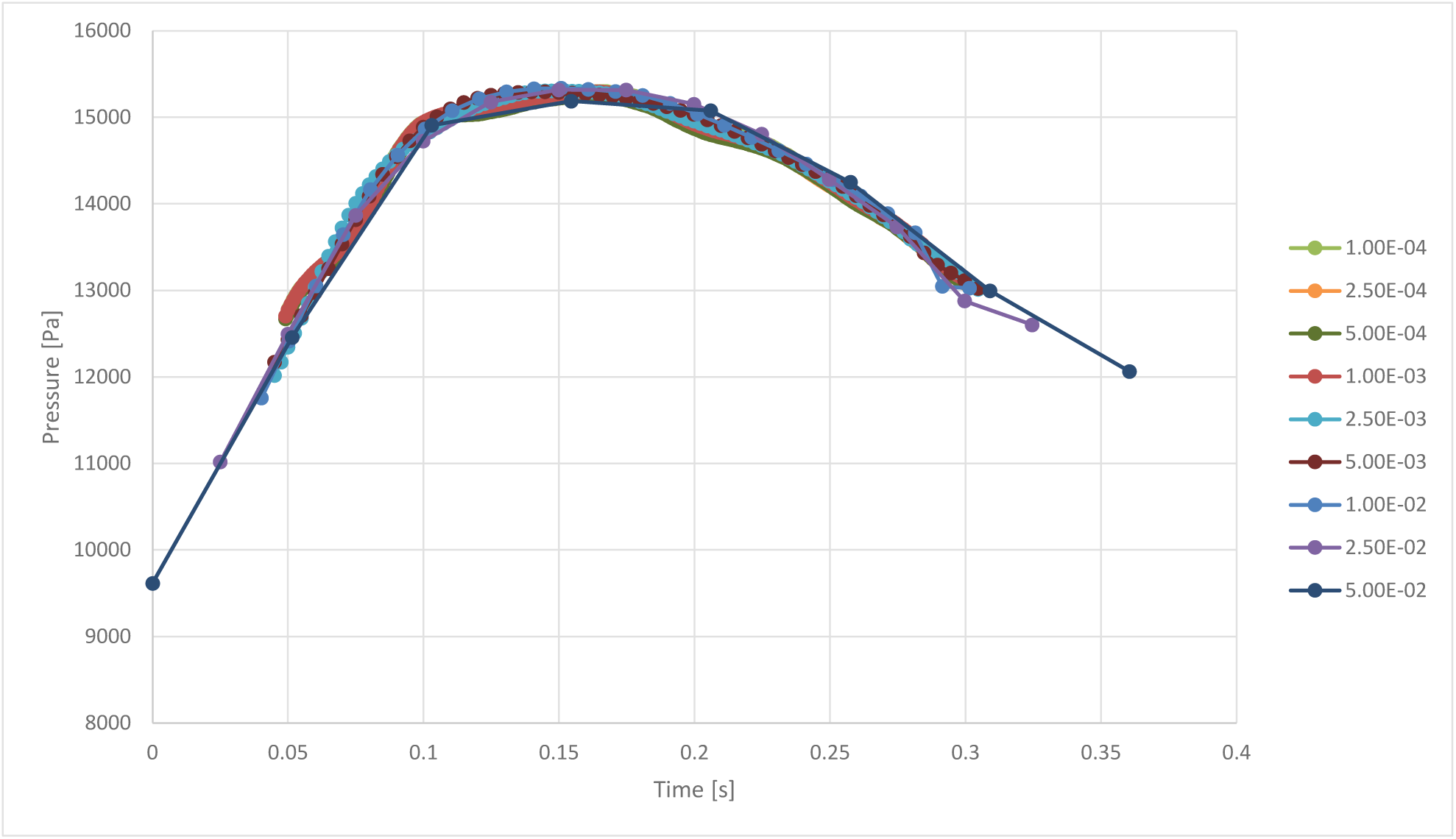

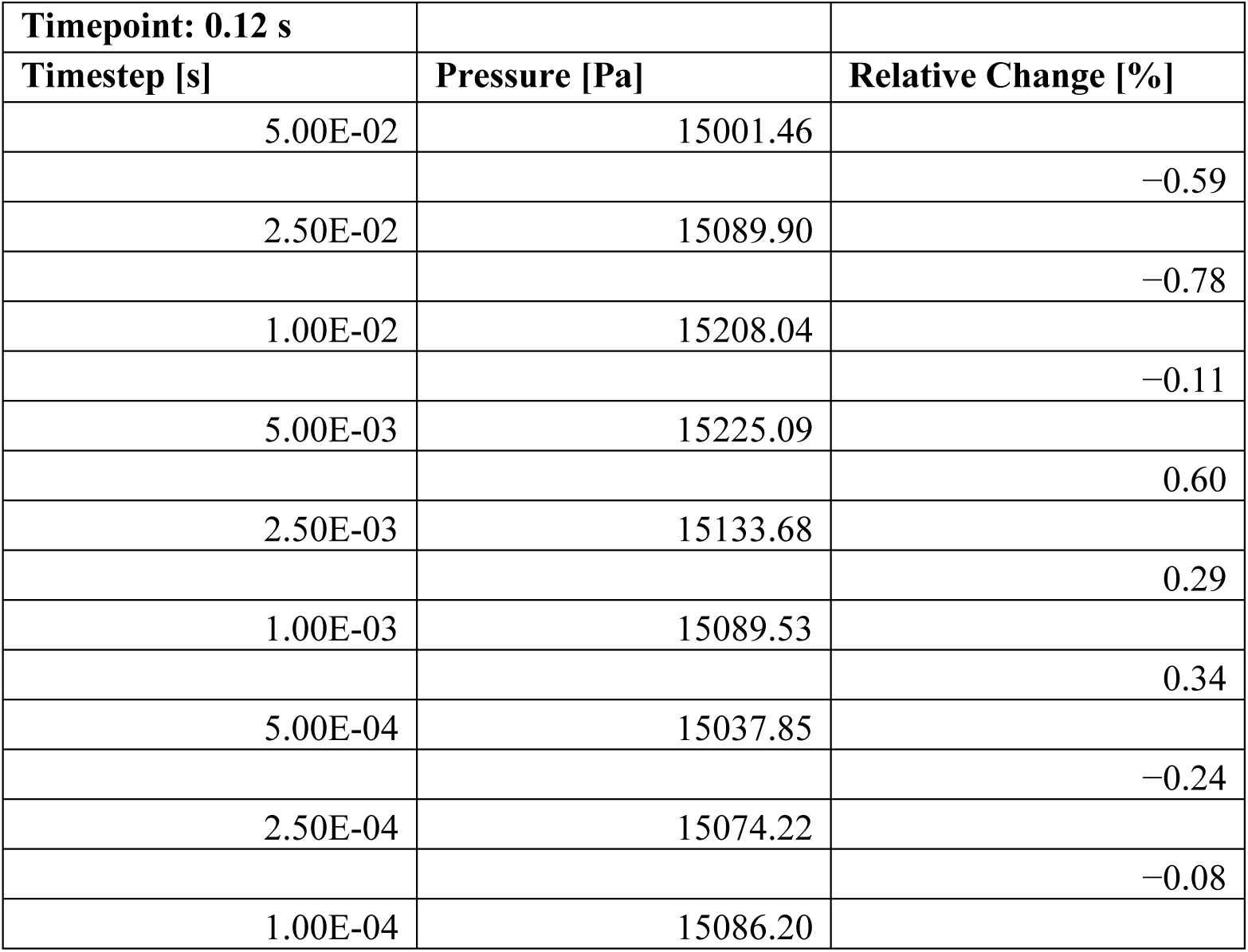

| Timepoint: 0.12 s |  |  |
| --- | --- | --- |
| Timestep [s] | Pressure [Pa] | Relative Change [%] |
| 5.00E-02 | 15001.46 |  |
|  |  | -0.59 |
| 2.50E-02 | 15089.90 |  |
|  |  | -0.78 |
| 1.00E-02 | 15208.04 |  |
|  |  | -0.11 |
| 5.00E-03 | 15225.09 |  |
|  |  | 0.60 |
| 2.50E-03 | 15133.68 |  |
|  |  | 0.29 |
| 1.00E-03 | 15089.53 |  |
|  |  | 0.34 |
| 5.00E-04 | 15037.85 |  |
|  |  | -0.24 |
| 2.50E-04 | 15074.22 |  |
|  |  | -0.08 |
| 1.00E-04 | 15086.20 |  |

#### Wall Shear Stress

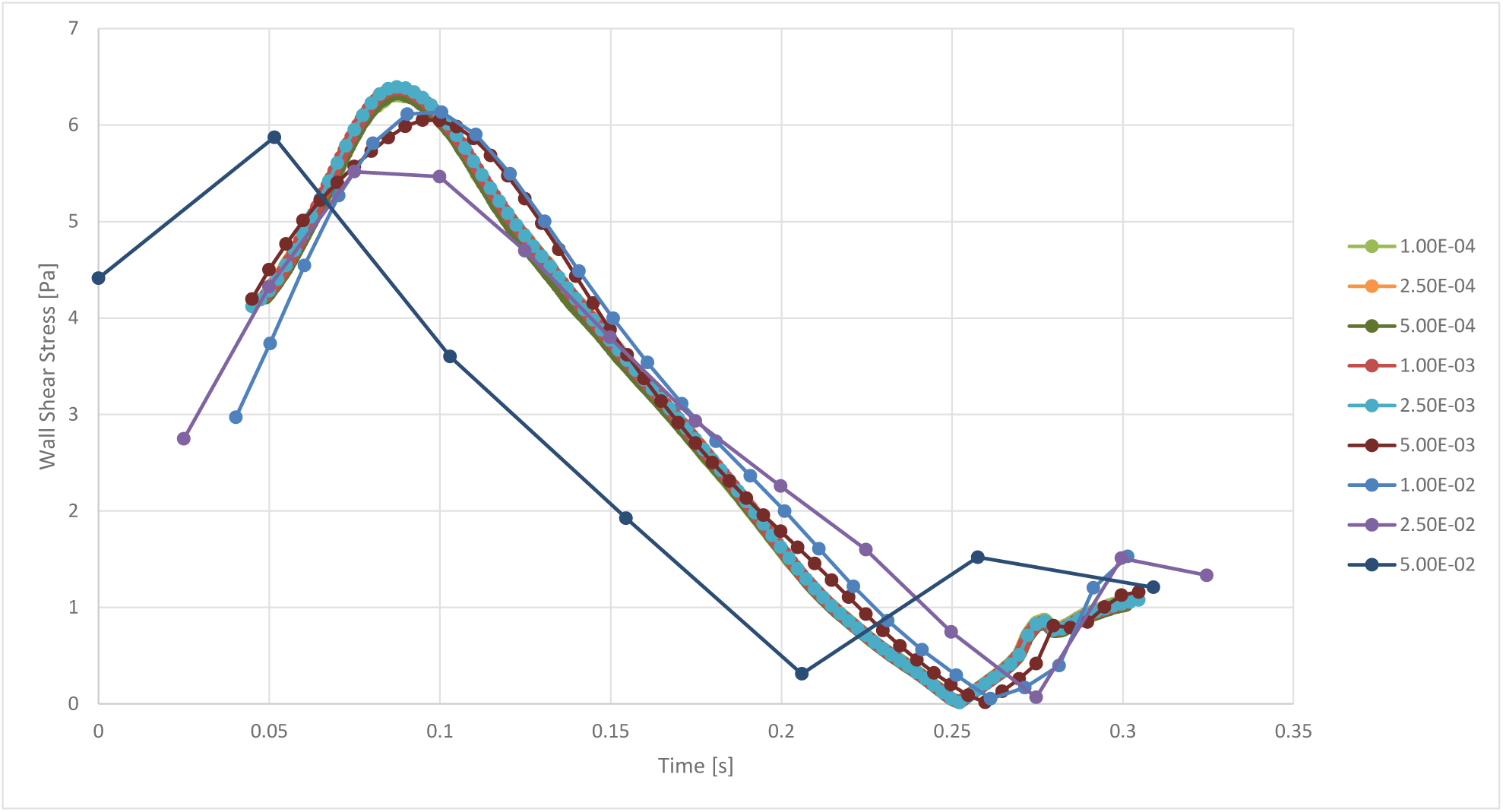

| Timepoint: 0.12 s |  |  |
| --- | --- | --- |
| Timestep [s] | Wall Shear Stress [Pa] | Relative Change [%] |
| 5.00E-02 | 5.122657593 |  |
|  |  | 5.69 |
| 2.50E-02 | 4.846970495 |  |
|  |  | -12.17 |
| 1.00E-02 | 5.518563629 |  |
|  |  | -1.91 |
| 5.00E-03 | 5.626055598 |  |
|  |  | 10.84 |
| 2.50E-03 | 5.075760048 |  |
|  |  | 0.25 |
| 1.00E-03 | 5.063169019 |  |
|  |  | -0.30 |
| 5.00E-04 | 4.988418528 |  |
|  |  | 0.81 |
| 2.50E-04 | 5.037535530 |  |
|  |  | -0.21 |
| 1.00E-04 | 5.048124887 |  |

## References

[1] D. and Loncar, C. Mathers, Projections of Global Mortality and Burden of Disease from 2002 to 2030, Plos Med. (2006). https://journals.plos.org/plosmedicine/article?id=10.1371/journal.pmed.0030442.

[2] R. Bauersachs, U. Zeymer, J.B. Brière, C. Marre, K. Bowrin, M. Huelsebeck, Burden of Coronary Artery Disease and Peripheral Artery Disease: A Literature Review, Cardiovasc. Ther. 2019 (2019). 10.1155/2019/8295054.

[3] C. Kasapis, H.S. Gurm, Current Approach to the Diagnosis and Treatment of Femoral-Popliteal Arterial Disease. A Systematic Review, Curr. Cardiol. Rev. 5 (2010) 296–311. 10.2174/157340309789317823.

[4] W.S. Aronow, Peripheral arterial disease in the elderly., Clin. Interv. Aging. 2 (2007) 645–654. 10.1161/01.atv.18.2.185.

[5] J. Shu, G. Santulli, Update on peripheral artery disease: Epidemiology and evidence-based facts, Atherosclerosis. 275 (2018) 379–381. 10.1016/j.atherosclerosis.2018.05.033.

[6] A.W. Bradbury, C.A. Moakes, M. Popplewell, L. Meecham, G.R. Bate, L. Kelly, I. Chetter, A. Diamantopoulos, A. Ganeshan, J. Hall, S. Hobbs, K. Houlind, H. Jarrett, S. Lockyer, J. Malmstedt, J. V. Patel, S. Patel, S.T. Rashid, A. Saratzis, G. Slinn, D.J.A. Scott, H. Zayed, J.J. Deeks, A vein bypass first versus a best endovascular treatment first revascularisation strategy for patients with chronic limb threatening ischaemia who required an infra-popliteal, with or without an additional more proximal infra-inguinal revascularisation pr, Lancet. 401 (2023) 1798–1809. 10.1016/S0140-6736(23)00462-2.

[7] A. Farber, M.T. Menard, M.S. Conte, J.A. Kaufman, R.J. Powell, N.K. Choudhry, T.H. Hamza, S.F. Assmann, M.A. Creager, M.J. Cziraky, M.D. Dake, M.R. Jaff, D. Reid, F.S. Siami, G. Sopko, C.J. White, M. van Over, M.B. Strong, M.F. Villarreal, M. McKean, E. Azene, A. Azarbal, A. Barleben, D.K. Chew, L.C. Clavijo, Y. Douville, L. Findeiss, N. Garg, W. Gasper, K.A. Giles, P.P. Goodney, B.M. Hawkins, C.R. Herman, J.A. Kalish, M.C. Koopmann, I.A. Laskowski, C. Mena-Hurtado, R. Motaganahalli, V.L. Rowe, A. Schanzer, P.A. Schneider, J.J. Siracuse, M. Venermo, K. Rosenfield, Surgery or Endovascular Therapy for Chronic Limb-Threatening Ischemia, N. Engl. J. Med. 387 (2022) 2305–2316. 10.1056/nejmoa2207899.

[8] P. Klinkert, A. Schepers, D.H.C. Burger, J.H. Van Bockel, P.J. Breslau, Vein versus polytetrafluoroethylene in above-knee femoropopliteal bypass grafting: Five-year results of a randomized controlled trial, J. Vasc. Surg. 37 (2003) 149–155. 10.1067/mva.2002.86.

[9] P. Klinkert, P.N. Post, P.J. Breslau, J.H. van Bockel, Saphenous vein versus PTFE for above-knee femoropopliteal bypass. A review of the literature, Eur. J. Vasc. Endovasc. Surg. 27 (2004) 357–362. 10.1016/j.ejvs.2003.12.027.

[10] S.F.C. Stewart, D.J. Lyman, Effects of an artery/vascular graft compliance mismatch on protein transport: A numerical study, Ann. Biomed. Eng. 32 (2004) 991–1006. 10.1023/B:ABME.0000032462.56207.65.

[11] M. Carrabba, P. Madeddu, Current Strategies for the Manufacture of Small Size Tissue Engineering Vascular Grafts, Front. Bioeng. Biotechnol. 6 (2018) 1–12. 10.3389/fbioe.2018.00041.

[12] B.H. Walpoth, G.L. Bowlin, The daunting quest for a small diameter vascular graft, Expert Rev. Med. Devices. 2 (2005) 647–651. 10.1586/17434440.2.6.647.

[13] E.M. Mahoney, K. Wang, H.H. Keo, S. Duval, K.G. Smolderen, D.J. Cohen, G. Steg, D.L. Bhatt, A.T. Hirsch, Vascular hospitalization rates and costs in patients with peripheral artery disease in the United States, Circ. Cardiovasc. Qual. Outcomes. 3 (2010) 642–651. 10.1161/CIRCOUTCOMES.109.930735.

[14] M.J.A. Lepäntalo, R. Houbballah, M. Raux, G. Lamuraglia, Lower extremity bypass vs endovascular therapy for young patients with symptomatic peripheral arterial disease, J. Vasc. Surg. 56 (2012) 545–554. 10.1016/j.jvs.2012.06.053.

[15] L. V. Thomas, V. Lekshmi, P.D. Nair, Tissue engineered vascular grafts - Preclinical aspects, Int. J. Cardiol. 167 (2013) 1091–1100. 10.1016/j.ijcard.2012.09.069.

[16] A. Bregy, S. Bogni, V.J.P. Bernau, I. Vajtai, F. Vollbach, A. Petri-Fink, M. Constantinescu, H. Hofmann, M. Frenz, M. Reinert, Solder doped polycaprolactone scaffold enables reproducible laser tissue soldering, Lasers Surg. Med. 40 (2008) 716–725. 10.1002/lsm.20710.

[17] M.S. Conte, M.J. Mann, H.F. Simosa, K.K. Rhynhart, R.C. Mulligan, Genetic interventions for vein bypass graft disease: A review, J. Vasc. Surg. 36 (2002) 1040–1052. 10.1067/mva.2002.129112.

[18] G. D. N., K. F., Coronary artery bypass grafting hemodynamics and anastomosis design: A biomedical engineering review, Biomed. Eng. Online. 12 (2013) 1–29. http://www.biomedical-engineering-online.com/content/12/1/129 http://ovidsp.ovid.com/ovidweb.cgi?T=JS&PAGE=reference&D=emed12&NEWS=N&AN=2014319326.

[19] V.S. Sottiurai, J.S.T. Yao, W.R. Flinn, R.C. Batson, Intimal hyperplasia and neointima: An ultrastructural analysis of thrombosed grafts in humans, Surgery. 93 (1983) 809–817.

[20] R.S. Taylor, R.J. McFarland, M.I. Cox, An investigation into the causes of failure of PTFE grafts, Eur. J. Vasc. Surg. 1 (1987) 335–343. 10.1016/S0950-821X(87)80061-0.

[21] L.M. Graham, H.S. Bassiouny, S.S. White, S.S. Glagov, E.S. Choi, D.P. Giddens, Anastomotic intimal hyperplasia: Mechanical injury or flow induced, J. Vasc. Surg. 15 (1992) 0708–0717. 10.1067/mva.1992.33849.

[22] A. Tiwari, K.S. Cheng, H. Salacinski, G. Hamilton, A.M. Seifalian, Improving the patency of vascular bypass grafts: The role of suture materials and surgical techniques on reducing anastomotic compliance mismatch, Eur. J. Vasc. Endovasc. Surg. 25 (2003) 287–295. 10.1053/ejvs.2002.1810.

[23] J.I. Brody, N.J. Pickering, Pathobiology of intimal hyperplasia, J. Vasc. Surg. 10 (1989) 583–585. 10.1016/0741-5214(89)90153-5.

[24] M.S. Lemson, J.H.M. Tordoir, M.J.A.P. Daemen, P.J.E.H.M. Kitslaar, Intimal hyperplasia in vascular grafts, Eur. J. Vasc. Endovasc. Surg. 19 (2000) 336–350. 10.1053/ejvs.1999.1040.

[25] Y. Jeong, Y. Yao, E.K.F. Yim, Current understanding of intimal hyperplasia and effect of compliance in synthetic small diameter vascular grafts, Biomater. Sci. 8 (2020) 4383–4395. 10.1039/d0bm00226g.

[26] A. Post, P. Diaz-Rodriguez, B. Balouch, S. Paulsen, S. Wu, J. Miller, M. Hahn, E. Cosgriff-Hernandez, Elucidating the role of graft compliance mismatch on intimal hyperplasia using an ex vivo organ culture model, Acta Biomater. 89 (2019) 84–94. 10.1016/j.actbio.2019.03.025.

[27] S.L. Meyerson, C.L. Skelly, M.A. Curi, U.M. Shakur, J.E. Vosicky, S. Glagov, L.B. Schwartz, The effects of extremely low shear stress on cellular proliferation and neointimal thickening in the failing bypass graft, J. Vasc. Surg. 34 (2001) 90–97. 10.1067/mva.2001.114819.

[28] S. Schoenborn, S. Pirola, M.A. Woodruff, M.C. Allenby, Fluid-Structure Interaction Within Models of Patient-Specific Arteries: Computational Simulations and Experimental Validations, IEEE Rev. Biomed. Eng. (2022). 10.1109/RBME.2022.3215678.

[29] L. Jia, L. Wang, F. Wei, H. Yu, H. Dong, B. Wang, Z. Lu, G. Sun, H. Chen, J. Meng, B. Li, R. Zhang, X. Bi, Z. Wang, H. Pang, A. Jiang, Effects of wall shear stress in venous neointimal hyperplasia of arteriovenous fistulae, Nephrology. 20 (2015) 335–342. 10.1111/nep.12394.

[30] W. Trubel, H. Schima, A. Moritz, F. Raderer, A. Windisch, R. Ullrich, U. Windberger, U. Losert, P. Polterauer, Compliance mismatch and formation of distal anastomotic intimal hyperplasia in externally stiffened and lumen-adapted venous grafts, Eur. J. Vasc. Endovasc. Surg. 10 (1995) 415–423. 10.1016/S1078-5884(05)80163-7.

[31] T.M.J. Van Bakel, C.J. Arthurs, F.J.H. Nauta, K.A. Eagle, J.A. Van Herwaarden, F.L. Moll, S. Trimarchi, H.J. Patel, C.A. Figueroa, Cardiac remodelling following thoracic endovascular aortic repair for descending aortic aneurysms, Eur. J. Cardio-Thoracic Surg. 55 (2019) 1061–1070. 10.1093/ejcts/ezy399.

[32] R. Jayendiran, B. Nour, A. Ruimi, Fluid-structure interaction (FSI) analysis of stent-graft for aortic endovascular aneurysm repair (EVAR): Material and structural considerations, J. Mech. Behav. Biomed. Mater. 87 (2018) 95–110. 10.1016/j.jmbbm.2018.07.020.

[33] R. Jayendiran, B. Nour, A. Ruimi, Computational fluid–structure interaction analysis of blood flow on patient-specific reconstructed aortic anatomy and aneurysm treatment with Dacron graft, J. Fluids Struct. 81 (2018) 693–711. 10.1016/j.jfluidstructs.2018.06.008.

[34] E. Tubaldi, M.P. Païdoussis, M. Amabili, Nonlinear dynamics of dacron aortic prostheses conveying pulsatile flow, J. Biomech. Eng. 140 (2018). 10.1115/1.4039284.

[35] S. Rahmani, A. Jarrahi, M. Navidbakhsh, M. Alizadeh, Investigating the performance of four specific types of material grafts and their effects on hemodynamic patterns as well as on von Mises stresses in a grafted three-layer aortic model using fluid-structure interaction analysis, J. Med. Eng. Technol. 41 (2017) 630–643. 10.1080/03091902.2017.1382590.

[36] G. E., X. A., G. G. S., Estimating the hemodynamic influence of variable main body-to-iliac limb length ratios in aortic endografts, Int. Angiol. 37 (2018) 41–45. http://www.embase.com/search/results?subaction=viewrecord&from=export&id=L620336614 10.23736/S0392-9590.17.03883-4.

[37] R. Ouwendijk, M. De Vries, P.M.T. Pattynama, M.R.H.M. Van Sambeek, M.W. De Haan, T. Stijnen, J.M.A. Van Engelshoven, M.G.M. Hunink, Imaging peripheral arterial disease: A randomized controlled trial comparing contrast-enhanced MR angiography and multi-detector row CT angiography, Radiology. 236 (2005) 1094–1103. 10.1148/radiol.2363041140.

[38] N. Byrne, M. Velasco Forte, A. Tandon, I. Valverde, T. Hussain, A systematic review of image segmentation methodology, used in the additive manufacture of patient-specific 3D printed models of the cardiovascular system, JRSM Cardiovasc. Dis. 5 (2016) 204800401664546. 10.1177/2048004016645467.

[39] J. Wang, P.K. Paritala, J.B. Mendieta, Y. Komori, O.C. Raffel, Y. Gu, Z. Li, Optical coherence tomography-based patient-specific coronary artery reconstruction and fluid–structure interaction simulation, Biomech. Model. Mechanobiol. 19 (2020) 7–20. 10.1007/s10237-019-01191-9.

[40] K.W. Beach, C.A. Isaac, D.J. Phillips, D.E. Strandness, An ultrasonic measurement of superficial femoral artery wall thickness, Ultrasound Med. Biol. 15 (1989) 723–728. 10.1016/0301-5629(89)90112-9.

[41] B.J. Lenz, H.C. Veldenz, J.W. Dennis, S. Khansarinia, L.R. Atteberry, M.H. Goldman, A three-year follow-up on standard versus thin wall ePTFE grafts for hemodialysis, J. Vasc. Surg. 28 (1998) 464–470. 10.1016/S0741-5214(98)70132-6.

[42] M.L. Marin, F.J. Veith, T.F. Panetta, R.E. Gordon, K.R. Wengerter, W.D. Suggs, L. Sanchez, M.K. Parides, Saphenous vein biopsy: A predictor of vein graft failure, J. Vasc. Surg. 18 (1993) 407–415. 10.1016/0741-5214(93)90258-N.

[43] X. Chen, J. Zhuang, H. Huang, Y. Wu, Fluid–structure interactions (FSI) based study of low-density lipoproteins (LDL) uptake in the left coronary artery, Sci. Rep. 11 (2021). 10.1038/s41598-021-84155-3.

[44] R. Campobasso, F. Condemi, M. Viallon, P. Croisille, S. Campisi, S. Avril, Evaluation of Peak Wall Stress in an Ascending Thoracic Aortic Aneurysm Using FSI Simulations: Effects of Aortic Stiffness and Peripheral Resistance, Cardiovasc. Eng. Technol. 9 (2018) 707–722. 10.1007/s13239-018-00385-z.

[45] M. Ghaffari, K. Tangen, A. Alaraj, X. Du, F.T. Charbel, A.A. Linninger, Large-scale subject-specific cerebral arterial tree modeling using automated parametric mesh generation for blood flow simulation, Comput. Biol. Med. 91 (2017) 353–365. 10.1016/j.compbiomed.2017.10.028.

[46] M.F. Rabbi, F.S. Laboni, M.T. Arafat, Computational analysis of the coronary artery hemodynamics with different anatomical variations, Informatics Med. Unlocked. 19 (2020). 10.1016/j.imu.2020.100314.

[47] J. Boyd, J.M. Buick, S. Green, Analysis of the Casson and Carreau-Yasuda non-Newtonian blood models in steady and oscillatory flows using the lattice Boltzmann method, Phys. Fluids. 19 (2007). 10.1063/1.2772250.

[48] P.H. Charlton, J.M. Harana, S. Vennin, Y. Li, P. Chowienczyk, J. Alastruey, Modeling arterial pulse waves in healthy aging: a database for in silico evaluation of hemodynamics and pulse wave indexes, Am. J. Physiol. - Hear. Circ. Physiol. 317 (2019) H1062–H1085. 10.1152/AJPHEART.00218.2019.

[49] Y. Ivanova, A. Yukhnev, L. Tikhomolova, E. Smirnov, A. Vrabiy, A. Suprunovich, A. Morozov, G. Khubulava, V. Vavilov, Experience of Patient-Specific CFD Simulation of Blood Flow in Proximal Anastomosis for Femoral-Popliteal Bypass, Fluids. 7 (2022). 10.3390/fluids7100314.

[50] F. Piatti, F. Sturla, M.M. Bissell, S. Pirola, M. Lombardi, I. Nesteruk, A. Della Corte, A.C. Alberto, E. Votta, 4D flow analysis of BAV-Related fluid-dynamic alterations: Evidences of wall shear stress alterations in absence of clinically-relevant aortic anatomical remodeling, Front. Physiol. 8 (2017). 10.3389/fphys.2017.00441.

[51] Z. Teng, S. Wang, A. Tokgoz, V. Taviani, J. Bird, U. Sadat, Y. Huang, A.J. Patterson, N. Figg, M.J. Graves, J.H. Gillard, Study on the association of wall shear stress and vessel structural stress with atherosclerosis: An experimental animal study, Atherosclerosis. 320 (2021) 38–46. 10.1016/j.atherosclerosis.2021.01.017.

[52] Z. Chen, H. Yu, Y. Shi, M. Zhu, Y. Wang, X. Hu, Y. Zhang, Y. Chang, M. Xu, W. Gao, Vascular Remodelling Relates to an Elevated Oscillatory Shear Index and Relative Residence Time in Spontaneously Hypertensive Rats, Sci. Rep. 7 (2017). 10.1038/s41598-017-01906-x.

[53] A. Karimi, M. Navidbakhsh, M. Alizadeh, A. Shojaei, A comparative study on the mechanical properties of the umbilical vein and umbilical artery under uniaxial loading, Artery Res. 8 (2014) 51–56. 10.1016/j.artres.2014.02.001.

[54] G.M. Bosi, B. Biffi, G. Biglino, V. Lintas, R. Jones, S. Tzamtzis, G. Burriesci, F. Migliavacca, S. Khambadkone, A.M. Taylor, S. Schievano, Can finite element models of ballooning procedures yield mechanical response of the cardiovascular site to overexpansion?, J. Biomech. 49 (2016) 2778–2784. 10.1016/j.jbiomech.2016.06.021.

[55] A. Schiavone, L.G. Zhao, A study of balloon type, system constraint and artery constitutive model used in finite element simulation of stent deployment, Mech. Adv. Mater. Mod. Process. 1 (2015). 10.1186/s40759-014-0002-x.

[56] M. Jadidi, S.A. Razian, M. Habibnezhad, E. Anttila, A. Kamenskiy, Mechanical, structural, and physiologic differences in human elastic and muscular arteries of different ages: Comparison of the descending thoracic aorta to the superficial femoral artery, Acta Biomater. 119 (2021) 268–283. 10.1016/j.actbio.2020.10.035.

[57] J. Lantz, J. Renner, M. Karlsson, Wall shear stress in a subject specific human aorta - Influence of fluid-structure interaction, Int. J. Appl. Mech. 3 (2011) 759–778. 10.1142/S1758825111001226.

[58] A. Caimi, M. Pasquali, F. Sturla, F.R. Pluchinotta, L. Giugno, M. Carminati, A. Redaelli, E. Votta, Prediction of post-stenting biomechanics in coarcted aortas: A pilot finite element study, J. Biomech. 105 (2020). 10.1016/j.jbiomech.2020.109796.

[59] S. Schoenborn, T. Lorenz, K. Kuo, D.F. Fletcher, M.A. Woodruff, S. Pirola, M.C. Allenby, Fluid-structure interactions of peripheral arteries using a coupled in silico and in vitro approach, Comput. Biol. Med. 165 (2023). 10.1016/j.compbiomed.2023.107474.

[60] F.A. Duck, Physical Properties of Tissue. A Comprehensive Reference Book, Med. Phys. 18 (1990) 834.

[61] C. Menichini, S. Pirola, B. Guo, W. Fu, Z. Dong, X.Y. Xu, High Wall Stress May Predict the Formation of Stent-Graft–Induced New Entries After Thoracic Endovascular Aortic Repair, J. Endovasc. Ther. 25 (2018) 571–577. 10.1177/1526602818791827.

[62] Suture Sizes and Suggested Indications for their use, Oxford Med. Educ. (2015) 2–3. https://oxfordmedicaleducation.com/surgery/suture-sizes-and-suggested-indications-for-their-use/.

[63] P.B. Dobrin, Some mechanical properties of polypropylene sutures: Relationship to the use of polypropylene in vascular surgery, J. Surg. Res. 45 (1988) 568–573. 10.1016/0022-4804(88)90146-1.

[64] Ineos Olefins & Polymers USA, Typical Engineering Properties of Polypropylene, Ineos Olefin. Polym. USA. (2010) 2. http://www.ineos.com/globalassets/ineos-group/businesses/ineos-olefins-and-polymers-usa/products/technical-information--patents/ineos-engineering-properties-of-pp.pdf.

[65] Poly(4-hydroxybutyrate), Polym. Database. (n.d.). https://polymerdatabase.com/polymers/Poly(4-hydroxybutyrate).html.

[66] D.P. Martin, S.F. Williams, Medical applications of poly-4-hydroxybutyrate: A strong flexible absorbable biomaterial, Biochem. Eng. J. 16 (2003) 97–105. 10.1016/S1369-703X(03)00040-8.

[67] G.N. Greaves, A.L. Greer, R.S. Lakes, T. Rouxel, Poisson’s ratio and modern materials, Nat. Mater. 10 (2011) 823–837. 10.1038/nmat3134.

[68] A.Y. Lee, H.C. Han, A nonlinear thin-wall model for vein buckling, Cardiovasc. Eng. Technol. 1 (2010) 282–289. 10.1007/s13239-010-0024-4.

[69] N.R. Tai, H.J. Salacinski, A. Edwards, G. Hamilton, A.M. Seifalian, Compliance properties of conduits used in vascular reconstruction, Br. J. Surg. 87 (2000) 1516–1524. 10.1046/j.1365-2168.2000.01566.x.

[70] A. Rassoli, N. Fatouraee, M. Shafigh, Uniaxial and Biaxial Mechanical Properties of the Human Saphenous Vein, Biomed. Eng. - Appl. Basis Commun. 27 (2015). 10.4015/S1016237215500507.

[71] C.S. Jorgensen, W.P. Paaske, Physical and mechanical properties of ePTFE stretch vascular grafts determined by time-resolved scanning acoustic microscopy, Eur. J. Vasc. Endovasc. Surg. 15 (1998) 416–422. 10.1016/S1078-5884(98)80203-7.

[72] M. Bouchet, M. Gauthier, M. Maire, A. Ajji, S. Lerouge, Towards compliant small-diameter vascular grafts: Predictive analytical model and experiments, Mater. Sci. Eng. C. 100 (2019) 715–723. 10.1016/j.msec.2019.03.023.

[73] S.K. Chimakurthi, S. Reuss, M. Tooley, S. Scampoli, ANSYS Workbench System Coupling: a state-of-the-art computational framework for analyzing multiphysics problems, Eng. Comput. 34 (2018) 385–411. 10.1007/s00366-017-0548-4.

[74] G.H. Lee, W. Heo, Y. Lee, T.H. Kim, H. Huh, S.W. Song, H. Ha, Fluid–structure interaction simulation of visceral perfusion and impact of different cannulation methods on aortic dissection, Sci. Rep. 13 (2023). 10.1038/s41598-023-27855-2.

[75] D. Shav, R. Gotlieb, U. Zaretsky, D. Elad, S. Einav, Wall shear stress effects on endothelial-endothelial and endothelial-smooth muscle cell interactions in tissue engineered models of the vascular wall, PLoS One. 9 (2014). 10.1371/journal.pone.0088304.

[76] S. Cheng, D. Fletcher, S. Hemley, M. Stoodley, L. Bilston, Effects of fluid structure interaction in a three dimensional model of the spinal subarachnoid space, J. Biomech. 47 (2014) 2826–2830. 10.1016/j.jbiomech.2014.04.027.

[77] T. Grus, L. Lambert, J. Matěcha, G. Grusová, M. Špaćek, M. Mlček, The ratio of diameters between the target artery and the bypass modifies hemodynamic parameters related to intimal hyperplasia in the distal end-to-side anastomosis, Physiol. Res. 65 (2016) 901–908. 10.33549/physiolres.933297.

[78] A. Desyatova, W. Poulson, J. MacTaggart, K. Maleckis, A. Kamenskiy, Cross-sectional pinching in human femoropopliteal arteries due to limb flexion, and stent design optimization for maximum cross-sectional opening and minimum intramural stresses, J. R. Soc. Interface. 15 (2018). 10.1098/rsif.2018.0475.

[79] C. M.J., L. X., L. W., Y. C., P. C.D., M. A., J. C.C., S. C., D. A., Therapeutic strategies to combat neointimal hyperplasia in vascular grafts, Expert Rev. Cardiovasc. Ther. 10 (2012) 635–647. http://www.embase.com/search/results?subaction=viewrecord&from=export&id=L364947399 http://dx.doi.org/10.1586/erc.12.33.

[80] G. Cao, X. Xuan, J. Hu, R. Zhang, H. Jin, H. Dong, How vascular smooth muscle cell phenotype switching contributes to vascular disease, Cell Commun. Signal. 20 (2022). 10.1186/s12964-022-00993-2.

[81] A.A. Owida, H. Do, Y.S. Morsi, Numerical analysis of coronary artery bypass grafts: An over view, Comput. Methods Programs Biomed. 108 (2012) 689–705. 10.1016/j.cmpb.2011.12.005.

[82] T. Meirson, E. Orion, C. Di Mario, C. Webb, N. Patel, K.M. Channon, Y. Ben Gal, D.P. Taggart, Flow patterns in externally stented saphenous vein grafts and development of intimal hyperplasia, J. Thorac. Cardiovasc. Surg. 150 (2015) 871–879. 10.1016/j.jtcvs.2015.04.061.

[83] R.S. Keynton, M.M. Evancho, R.L. Sims, N. V. Rodway, A. Gobin, S.E. Rittgers, Intimal hyperplasia and wall shear in arterial bypass graft distal anastomoses: An in vivo model study, J. Biomech. Eng. 123 (2001) 464–473. 10.1115/1.1389461.

[84] F. Migliavacca, G. Dubini, Computational modeling of vascular anastomoses, Biomech. Model. Mechanobiol. 3 (2005) 235–250. 10.1007/s10237-005-0070-2.

[85] I.J. Freshwater, Y.S. Morsi, T. Lai, The effect of angle on wall shear stresses in a LIMA to LAD anastomosis: Numerical modelling of pulsatile flow, Proc. Inst. Mech. Eng. Part H J. Eng. Med. 220 (2006) 743–757. 10.1243/09544119JEIM126.

[86] M. Sankaranarayanan, L.P. Chua, D.N. Ghista, Y.S. Tan, Computational model of blood flow in the aorto-coronary bypass graft, Biomed. Eng. Online. 4 (2005). 10.1186/1475-925X-4-14.

[87] T. Cao, Z. Jiang, H. Zhao, K.Q. Zhang, K. Meng, Numerical simulation to study the impact of compliance mismatch between artificial and host blood vessel on hemodynamics, Med. Nov. Technol. Devices. 15 (2022). 10.1016/j.medntd.2022.100152.

[88] G. Yao, H. Li, X. Zuo, C. Wang, Y. Xiao, Y. Zhao, X. Wang, Oscillatory shear stress promotes vein graft intimal hyperplasia via NADPH oxidase-related pathways, Front. Surg. 10 (2023). 10.3389/fsurg.2023.1073557.

[89] Y.M. Kim, S.J. Kim, R. Tatsunami, H. Yamamura, T. Fukai, M.U. Fukai, ROS-induced ROS release orchestrated by Nox4, Nox2, and mitochondria in VEGF signaling and angiogenesis, Am. J. Physiol. - Cell Physiol. 312 (2017) C749–C764. 10.1152/ajpcell.00346.2016.

[90] Y. Qiu, J.M. Tarbell, Computational simulation of flow in the end-to-end anastomosis of a rigid graft and a compliant artery, ASAIO J. 42 (1996). 10.1097/00002480-199609000-00078.

[91] A. Leuprecht, K. Perktold, M. Prosi, T. Berk, W. Trubel, H. Schima, Numerical study of hemodynamics and wall mechanics in distal end-to-side anastomoses of bypass grafts, J. Biomech. 35 (2002) 225–236. 10.1016/S0021-9290(01)00194-4.

